# Unsupervised transcriptomic analysis of paired pre- and post-treatment specimens reveals divergent chemoimmunomodulatory induction trajectories in breast cancer

**DOI:** 10.64898/2026.09.24.754178

**Authors:** Iasmim Lopes de Lima, Elizabeth Molchan, Kathleen H. Streeks, Kennedy L. Coleman, Mariana S. Makarem, Nathan Seligson, Mohammed O. Gbadamosi

**Affiliations:** Department of Pharmacotherapy and Translational Research, College of Pharmacy, University of Florida, Gainesville, Florida; Brain Tumor Immunotherapy Program, Preston A. Wells Jr. Center for Brain Tumor Therapy, University of Florida, Gainesville, Florida; University of Florida Health Cancer Center, Gainesville, Florida

**Keywords:** chemotherapy, chemoimmunomodulation, immunotherapy, immune-checkpoint inhibitors, breast cancer

## Abstract

The immunomodulatory effects of chemotherapy (chemoimmunomodulation; CIM) are clinically consequential and heterogeneous, yet no systematic framework exists for classifying the immunomodulatory trajectory a tumor follows in response to treatment (CIM trajectory). Here, we present the CIM Induction Classifier (CIMIC), an unsupervised clustering pipeline leveraging delta gene expression across 3,189 CIM-related genes to classify specimens’ chemoimmunomodulatory trajectory. Applied to two pre- and post-chemotherapy breast cancer (BC) datasets (NKI/SMC, N = 36; NEO, N = 19) and nine epirubicin-perturbed triple-negative BC (TNBC) cell lines, CIMIC identified two divergent CIM trajectories: a functional CIM (Fun-CIM) trajectory, broadly conserved across tumors and cell lines and characterized by induction of inflammatory cell death, antigen presentation, viral mimicry, and adaptive immune activation programs, and a dysfunctional CIM (Dys-CIM) trajectory, characterized by induction of proteostatic and metabolic stress-adaptation programs, reduced immune cell abundances and cytotoxic activity, and enrichment of aggressive BC subtypes. Using survival and longitudinal transcriptomic data in NKI/SMC (N = 20), treatment-induced increases in Fun-CIM-associated genes and ssGSEA scores were associated with reduced recurrence, whereas Dys-CIM-associated genes and scores were associated with increased recurrence. In multivariable analyses within independent chemotherapy-treated BC cohorts (METABRIC, N = 412; SCAN-B, N = 2,462), higher baseline Fun-CIM ssGSEA scores were associated with better outcomes, whereas higher baseline Dys-CIM ssGSEA scores were associated with worse outcomes. These findings establish CIM as a dynamic, trajectory-level process and position CIMIC as a framework for defining CIM trajectories and supporting future efforts to identify predictors, mechanisms, and therapeutic strategies that maximize beneficial CIM.

## 1. Introduction

Despite advancements in breast cancer (BC) therapy, chemotherapy remains the predominant front-line pharmacologic intervention and a cornerstone of curative treatment regimens (1). However, its clinical efficacy and effects on the tumor microenvironment (TME) are highly variable (2). A key contributor to this observed heterogeneity is chemoimmunomodulation (CIM), which comprises the tumor-intrinsic and extrinsic shifts in tumor immunity elicited by chemotherapy (3,4). Previous studies have demonstrated that CIM induces a diversity of compositional and functional changes that vary with treatment regimen and molecular context (3,4).

Specifically, chemotherapy can induce both immunostimulatory and immunosuppressive remodeling of the TME. For instance, some chemotherapeutics can induce immunogenic cell death (ICD), characterized by the release or exposure of damage-associated molecular patterns (DAMPs), type I interferon (IFN) signaling, and the upregulation of antigen presentation programs, which collectively drive TME immune activation and tumor-directed cytotoxicity (4–9). Engagement of distinct regulated cell death (RCD) pathways, such as necroptosis, pyroptosis, PANoptosis, and others, further influences immunogenic potential by promoting robust immunostimulatory effects within the TME (10–14). Beyond cell death-associated mechanisms, numerous agents exert dose- and class-specific effects on immune cell abundance and functional state, orchestrating immunostimulatory CIM that occurs in tandem with or independent of ICD or RCD modalities (6). These properties have motivated efforts to exploit the immunomodulatory effects of chemotherapy through optimized treatment strategies and rational combinations with immunotherapy (15,16). In contrast, chemotherapeutic agents can also promote immunosuppressive CIM effects, including the recruitment and expansion of protumoral myeloid-derived suppressor cells and regulatory immune subsets and activation of tolerogenic cytokine signaling, dampening antitumor immunity (17,18). Indeed, RCD pathways, like ferroptosis, act as a double-edged sword, carrying both immunostimulatory and immunosuppressive consequences, depending on the context (19).

Fundamentally, CIM is an inherently dynamic process encompassing the evolution of tumor-intrinsic stress responses and TME remodeling over the course of treatment (4,20). Consequently, evaluating individual immunomodulatory processes or molecular profiling at static pre- and post-treatment timepoints cannot comprehensively reflect CIM and fails to capture how evolving signals resolve into the global net immunomodulatory trajectory a tumor follows in response to therapy (CIM trajectory). Furthermore, longitudinal profiling of patients with BC treated with neoadjuvant chemotherapy has demonstrated that chemoimmunomodulatory responses evolve over the entire course of therapy and differ across BC subtypes and treatment responses (21), suggesting that CIM, and its downstream functional and clinical consequences, may vary by baseline tumor characteristics and treatment regimen, dose, and schedule (16,21). Thus, CIM represents a heterogeneous yet clinically relevant process requiring systematic investigation using a comprehensive, trajectory-based approach (3,4,21–23).

However, despite these insights, CIM remains critically understudied. Of studies investigating CIM, most have focused on individual immunomodulatory processes or relied on static molecular profiling at pre- and post-treatment timepoints, rather than analyzing the CIM trajectory from a global perspective. This limited focus likely reflects the absence of a robust, comprehensive classification framework for capturing CIM trajectories at a global level and reliably distinguishing immunostimulatory, beneficial CIM trajectories from immunosuppressive, detrimental CIM trajectories. This knowledge gap has limited our ability to identify the biological characteristics that define and drive divergent CIM trajectories and has impeded efforts to determine which therapeutic strategies in which contexts yield maximum immunomodulatory benefit.

To address this challenge, we developed the CIM Induction Classifier (CIMIC), a novel iterative unsupervised clustering pipeline that identifies distinct, population-level CIM trajectories by clustering individual tumors according to similarities in their chemotherapy-induced transcriptional remodeling. The CIMIC pipeline uses delta (Δ) gene expression (GE) values for 3,189 genes spanning 19 CIM-related pathways, calculated from paired pre- and post-treatment tumor transcriptomic data to classify tumors into distinct and stable CIM trajectories. Here, we apply the CIMIC pipeline to pre- and post-treatment data from BC patients and triple-negative breast cancer (TNBC) cell lines and investigate the transcriptomic, immunologic, and clinical characteristics associated with divergent CIM trajectories. Our pipeline and findings provide a new framework for defining the immunomodulatory trajectory of tumor remodeling following chemotherapy, which may inform future efforts to better characterize, understand, predict, and therapeutically optimize CIM to maximize clinical benefit.

## 2. Methods

### Patient Cohorts

Two datasets of pre- and post-treatment RNA-seq data from BC patient tumors were utilized in this study.

The first dataset (NKI/SMC) consisted of data from two studies (GSE191127; NKI and GSE123845; SMC). The NKI study contained data from 20 patients with locally advanced BC treated with neoadjuvant chemotherapy (NACT) in clinical trials (NCT00448266, NCT01057069) at the Netherlands Cancer Institute (24). Data from matched pre-treatment core biopsies and post-treatment surgical specimens were obtained for this study. No patient achieved pathological complete response (pCR) after NACT, and all pre- and post-treatment specimens contained > 50% tumor cells. Given that patient specimens at each time point were collected in duplicate, average GE values across the technical duplicates were computed for each patient and used for downstream analyses (24). For the NKI cohort, patient-level clinical follow-up data including recurrence-free survival (RFS) and overall survival (OS) were available for a subset of patients (N = 20; 6 events) and were used for survival analyses. Survival data for the SMC cohort were not available. The SMC study contained data from 16 selected patients with locally advanced (n = 15) and metastatic (n = 1) BC treated with NACT in the clinical trial (NCT02591966) at Samsung Medical Center (SMC). Data from matched pre-treatment biopsies and post-treatment surgical specimens collected following approximately 6 months of therapy were obtained for this study. None of the selected patients achieved pCR, and the specimens contained a mean tumor purity of 62.6%. All patients across both cohorts received standard anthracycline-, taxane-, and/or cyclophosphamide-based regimens. Data from NKI and SMC were harmonized for analyses. Details on harmonization of transcriptomic data are described in **Supplemental Methods SM1,** and patient demographics data are shown in **Tables 1** and **2** for the NKI and SMC cohorts, respectively (21).

The second dataset (GSE122630; NEO) included 19 selected patients with primary operable, grade II-III, invasive BC treated with neoadjuvant chemotherapy at the Edinburgh Breast Unit at the Western General Hospital. This data was used to validate CIMIC’s ability to identify divergent CIM trajectories. We focused our analyses on patients with matched pre-treatment core biopsies and surgical resection specimens who were HER2-negative (N = 19), thereby focusing on patients who had undergone TME remodeling in response to cytotoxic chemotherapy and limiting the impact of targeted agents like trastuzumab on the analysis. Patients were treated with standard neoadjuvant chemotherapy regimens, primarily consisting of three cycles of fluorouracil (5-FU), epirubicin, and cyclophosphamide followed by docetaxel. Transcriptomic profiling was performed on fresh frozen samples using the Ion AmpliSeq Transcriptome Human Gene Expression Kit, as described previously (25). Patient demographic data are shown in **Supplemental Table 1**.

Survival analyses assessing the association between pre-treatment GE and clinical outcomes were performed using independent BC cohorts, including the Taxonomy of Breast Cancer International Consortium (METABRIC) and the Sweden Cancerome Analysis Network - Breast (SCAN-B) datasets. The METABRIC cohort comprises log₂-transformed microarray-based gene expression data from fresh-frozen primary BC specimens. Corresponding clinical annotations were obtained from the cBioPortal for Cancer Genomics (26). The SCAN-B cohort is a prospective, population-based BC research program initiated in 2010 across southern Sweden. Gene expression data quantified as fragments per kilobase of transcript per million mapped reads (FPKM), along with associated clinical characteristics, were downloaded from the Mendeley Data repository (27). SCAN-B expression values were log-transformed using log₂ (FPKM + 0.1) prior to analysis. Only patients who received systemic chemotherapy within either cohort were included in the downstream analysis (METABRIC, *n =* 412; SCAN-B, *n =* 2,462). Patient demographics data are shown in **Table 3** for the METABRIC and SCAN-B cohorts.

### Development of the Chemoimmunomodulation Induction Classifier (CIMIC)

The chemoimmunomodulation induction classifier (CIMIC) is an unsupervised stability-guided, iterative computational pipeline developed to identify distinct population-level CIM trajectories by clustering individual tumor specimens based on their treatment-induced transcriptional remodeling (**Figure 1**). CIMIC uses within-specimen changes in gene expression calculated as ΔGE = log2[TPM_post-treatment_ + 1] - log2[TPM_pre-treatment_ + 1]) across 3,189 genes spanning 19 CIM-related pathways (**Supplemental Table 2**) (4,28,29). By clustering tumors based on their treatment-induced transcriptional changes, CIMIC-identified CIM trajectories are based on similarities in the direction and magnitude of chemotherapy-induced transcriptional remodeling rather than differences in absolute pre- or post-treatment expression. CIM-related pathways were selected to encompass immunostimulatory and immunosuppressive, tumor-intrinsic and tumor-extrinsic CIM programs (4,30). Detailed pathway selection criteria are provided in **Supplemental Methods SM2**.

**Figure 1.**
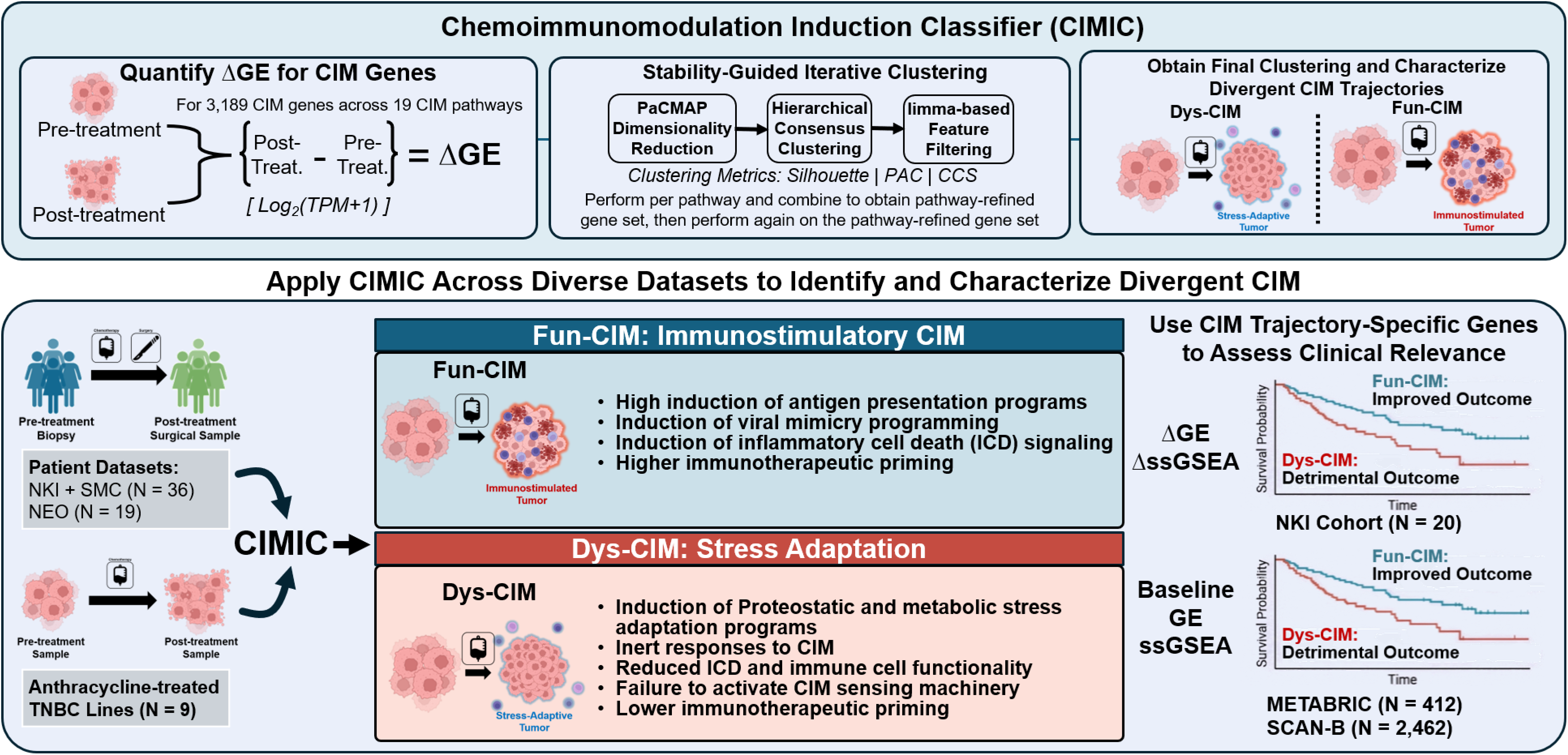
Study design and overview of the Chemoimmunomodulation Induction Classifier (CIMIC). CIMIC quantifies within-patient transcriptional changes as delta gene expression (ΔGE; post-treatment log₂[TPM + 1] − pre-treatment log₂[TPM + 1]) across 3,189 genes representing 19 chemoimmunomodulatory (CIM) pathways. Pathway-level PaCMAP dimensionality reduction, hierarchical consensus clustering, and limma-based feature filtering are iteratively applied to identify robust, biologically distinct CIM trajectories. CIMIC was applied to matched pre-treatment biopsy and post-treatment surgical specimens from patients with breast cancer who received neoadjuvant chemotherapy in the combined NKI/SMC cohort (N = 36) and the independent NEO validation cohort (N = 19), as well as to nine triple-negative breast cancer (TNBC) cell lines treated with epirubicin. Across datasets, two divergent transcriptional chemoimmunomodulation trajectories were identified. Functional CIM (Fun-CIM) was characterized by induction of antigen-presentation programs, viral-mimicry signaling, immunogenic cell-death (ICD) pathways, and greater immunotherapeutic priming. Dysfunctional CIM (Dys-CIM) was characterized by induction of proteostatic and metabolic stress-adaptation programs, reduced ICD signaling and immune-cell functionality, failure to activate CIM-sensing machinery, and limited immunotherapeutic priming. The clinical relevance of trajectory-associated transcriptional programs was evaluated using longitudinal changes in single-sample gene set enrichment analysis (ssGSEA) scores in the NKI cohort with available survival follow-up (N = 20) and baseline ssGSEA scores in independent chemotherapy-treated METABRIC (N = 412) and SCAN-B (N = 2,462) cohorts. CIM, chemoimmunomodulation; ΔGE, change in gene expression; PaCMAP, Pairwise Controlled Manifold Approximation; ssGSEA, single-sample gene set enrichment analysis; TNBC, triple-negative breast cancer.

The CIMIC pipeline first performs within-pathway refinement for each of the 19 CIM-related pathways. For each pathway, samples are clustered by subjecting their ΔGE values to dimensionality reduction using PaCMAP (v0.8.0) (31), followed by hierarchical consensus clustering implemented in the ConsensusClusterPlus package (v1.73.0) (32), using Euclidean distance and Ward’s minimum-variance linkage. The optimal number of clusters (*k)* used for consensus clustering procedure is determined using a composite ranking framework that integrates measures of cluster separation and stability metrics across the tested range of *k* values. After clustering at the optimal *k*, limma-based linear models with empirical-Bayes variance moderation (v3.66.0) (33) are used to identify nondiscriminatory genes based on false discovery rate (FDR)-adjusted p values (p_adj_ > 0.05). This optimal k determination, clustering, and feature filtering procedure is iteratively repeated for each pathway until convergence, defined as the point at which no additional genes are excluded in a subsequent clustering and feature-filtering iteration.

Once within-pathway refinement has been performed for all 19 CIM-related pathways, the final discriminatory genes retained within each pathway are combined to generate a pathway-refined global gene set. The same iterative, stability-guided refinement process is then applied to the pathway-refined global gene set to further remove redundant and non-discriminatory genes across pathways, with cluster assignments at the convergence of this procedure reflecting the assignment of tumors to CIMIC-identified, population-level CIM trajectories. Details on the iterative, stability-guided refinement process and other dimensions of CIMIC are further described in **Supplemental Methods SM3**.

### In vitro triple-negative breast cancer cell line drug treatment and RNA-seq for orthogonal validation

Early passages of nine TNBC cell lines (ATCC, VA, USA) were cultured under recommended conditions. Epirubicin (Selleckchem, TX, USA) Drug sensitivity for each cell line was assessed using a 9-point dose-response curve generated using the CellTiter-Glo assay (CTG; Promega, WI, US) over the course of 48 hours. IC_30_ values were determined per cell line by fitting dose-response data to a four-parameter logistic model and conducting extrapolation. For RNA-sequencing, 5x10^5^ cells were subjected to 48-hour treatment with DMSO, representing a pre-treatment control, or epirubicin at the IC_30_ value derived for that cell line. Following treatment, cells were collected, and total RNA was extracted using the RNeasy Mini Plus Kit (QIAGEN, MD, USA) following the manufacturer’s instructions. RNA-sequencing libraries were prepared using the KAPA HyperPrep mRNA kit (Roche, IN, USA) and paired-end sequencing was performed on the NovaSeq 6000 (Signios Bio, CA, USA). High-quality reads were then aligned to GRCh38 and quantified at the gene level; expression was converted to TPM and log2-transformed as log2(TPM+1) for analysis, mirroring the patient pipeline. For each gene, ΔGE (post-treatment log2[TPM + 1] - pre-treatment log2 (TPM + 1]) values were calculated by subtracting the average pre-treatment GE values from post-treatment value across biological replicates of each cell line and utilized for the CIMIC pipeline. The CIMIC pipeline was restricted to tumor-intrinsic CIM-related pathways activated for cell line analyses. In addition, the duplicateCorrelation function was incorporated into the limma modeling framework within CIMIC and downstream analyses to account for within-cell line correlation among biological replicates, enabling CIM trajectories to reflect independent cell line effects while retaining information from repeated measurements.

### Statistical Analyses

Wilcoxon rank sum and Wilcoxon signed-rank tests were used for unpaired and paired comparisons, respectively, and chi-square or Fisher’s exact tests were used to examine the differences between categories based on whether data met test assumptions. Cluster-defining genes were identified using a point-biserial correlation values (r_pb_) derived from limma’s moderated *t*-statistics, quantifying the strength and direction of association between gene-level ΔGE and CIM trajectory (34). Enrichment analyses were performed using clusterprofiler (v4.12.6) (35,36). The fry function within limma was used to assess coordinated changes within the CIM-related pathways and other predefined gene sets between groups. Immune cell type deconvolution was performed using CIBERSORTx with the LM22 signature matrix, employing 1,000 permutations and B-mode batch correction (https://cibersortx.stanford.edu/) (37) or the immunedeconv R package (v2.1.0), integrating multiple algorithms (quanTIseq, EPIC, xCell, MCP-counter, ESTIMATE, TIMER, and ConsensusTME) (38). Paired biopsy-to-surgery changes within CIM clusters were evaluated using Δ values (surgery − biopsy). For the NKI and NEO cohorts, the PAM50 molecular subtypes were predicted using Genefu package (39). Baseline transcriptional differences between Fun-CIM and Dys-CIM tumors were evaluated using limma. Gene-wise linear models were fitted using lmFit function within limma with CIM group as the explanatory variable (expression ∼ CIM_group). Moderated statistics were calculated using eBayes function within limma. Sample-level Fun-CIM and Dys-CIM enrichment scores were calculated using single-sample gene set enrichment (ssGSEA) analysis implemented in the GSVA package v2.4.9. Associations between individual genes and ssGSEA scores were evaluated using univariate and multivariable Cox proportional hazards models and, where applicable, Kaplan-Meier analyses, employing the survminer (v0.5.1) and survival (v3.8.3) packages (40,41). The association of treatment-induced ΔGE and ssGSEA scores was evaluated with RFS and OS using data from the NKI cohort, whereas the association between baseline ssGSEA scores and clinical outcomes was evaluated in chemotherapy-treated patients from the METABRIC (OS, DSS, and RFI) and SCAN-B (OS, DRFi, and RFI) cohorts. Cross-dataset conservation of CIM trajectory-associated genes was assessed using Stouffer’s method (42) to combine p values from directionally concordant gene-level and program-level analyses across NKI/SMC and TNBC cell line datasets or NKI/SMC, NEO, and TNBC cell line datasets, as indicated. Genes showing significant combined evidence were subsequently subjected to overrepresentation analysis to identify biological programs conserved across Fun-CIM tumors and cell lines. Where indicated, Z-score normalization was used for visualization. In cases of multiple testing, p values were FDR-adjusted, unless otherwise specified (43). All analyses were performed using the R platform (v4.5.0) unless otherwise specified.

## 3. Results

### CIMIC Identifies Distinct, Stable CIM trajectories

The overall study design is shown in **Figure 1**. To define population-level CIM trajectories, we applied the CIMIC to ΔGE data derived from paired pre- and post-treatment tumor transcriptomes from the NKI/SMC dataset (N = 36). CIMIC identified *k* = 2 as the optimal clustering solution according to our composite ranking framework. Specifically, the optimal cluster solution exhibited moderate overall silhouette separation (*s_avg_*= 0.569), no assignment ambiguity (PAC = 0), and maximal cluster stability (cluster consensus score = 1.00) (**Supplemental Table 3**). PaCMAP projections demonstrated the segregation of tumors into two major clusters (**Figure 2A**). Based on subsequent transcriptomic characterization (described below), these clusters were designated as a functional CIM trajectory (C2; Fun-CIM, n = 25), which was immune-activating, and a dysfunctional CIM trajectory (C1; Dys-CIM, n = 11), which was stress-adaptive and immunosuppressive. Heatmap visualization of ΔGE across all protein-coding genes revealed distinct, opposing transcriptional response to chemotherapy between Fun-CIM and Dys-CIM tumors, indicating transcriptomic distinction (**Figure 2B**).

**Figure 2.**
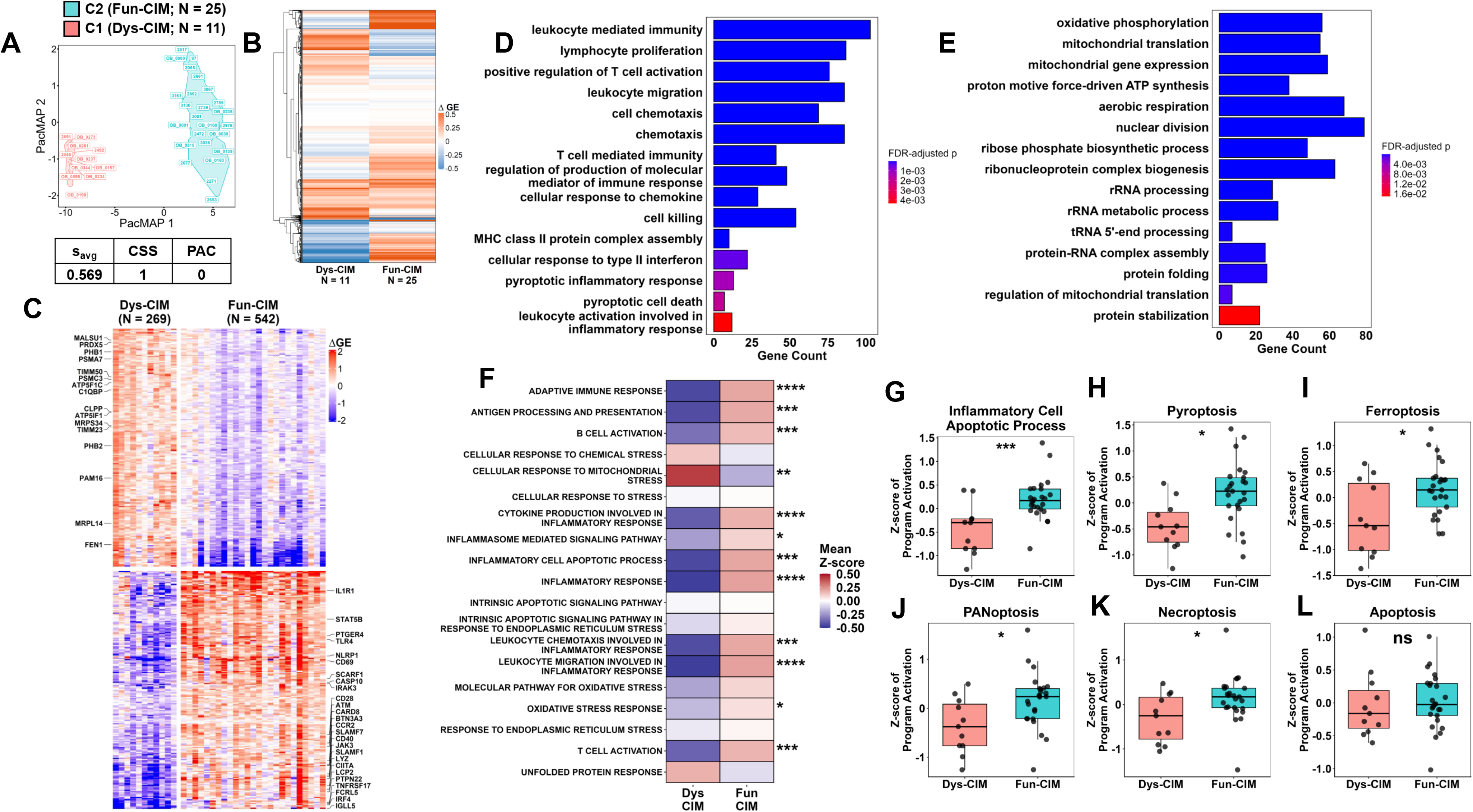
CIMIC identifies transcriptionally distinct chemoimmunomodulation induction trajectories in breast cancer tumors. **(A)** PaCMAP projection showing the segregation of tumors from the combined NKI/SMC cohort into two CIM trajectories: Dysfunctional CIM (C1, Dys-CIM; N = 11) and Functional CIM (C2, Fun-CIM; N = 25). Average silhouette score from ΔGE and PaCMAP reduced space (s_avg_), cluster consensus score (CSS), and proportion of ambiguous clustering (PAC) metrics are indicated. **(B)** Heatmap illustrating coordinated global transcriptomic differences in ΔGE (post-treatment log2[TPM + 1] - pre-treatment log2[TPM + 1]) in Fun-CIM and Dys-CIM tumors following chemotherapy**. (C)** Heatmap of the top 200 CIM-trajectory associated genes (|r_pb_| ≥ 0.6; FDR-adjusted p ≤ 0.05) for Fun-CIM and Dys-CIM trajectories. **(D-E)** Top enriched Gene Ontology (GO) biological processes among genes preferentially induced in Fun-CIM **(D)** and Dys-CIM **(E)** tumors following chemotherapy (FDR-adjusted p < 0.15). **(F)** Heatmap showing the relative induction of selected chemoimmunomodulatory pathways used for clustering, expressed as the Z-scored average ΔGE, in Fun-CIM and Dys-CIM tumors. **(G-H)** Boxplots comparing the relative differential induction (Z-score of average ΔGE values) of cell-death programs between Fun-CIM and Dys-CIM tumors, including inflammatory cell apoptotic process **(G)**, pyroptosis **(H)**, ferroptosis **(I)**, PANoptosis **(J)**, necroptosis **(K)**, and apoptosis **(L)**. Fun-CIM tumors exhibited significantly greater induction of inflammatory cell apoptotic, pyroptotic, ferroptotic, PANoptotic, and necroptotic programs, whereas induction of apoptosis did not significantly differ between trajectories. FDR-adjusted *p* values are indicated as follows: * ≤ 0.05; ** ≤ 0.01; *** ≤ 0.005; **** ≤ 0.001; ns, not significant.

### CIMIC-Identified CIM trajectories Comprise Induction of Divergent Immune and Tumor-Intrinsic Transcriptional Programs

Point-biserial correlations derived from limma’s moderated *t*-statistics (r_pb_) between Fun-CIM and Dys-CIM tumors identified 811 differentially induced genes, with 542 genes associated with Fun-CIM and 269 associated with Dys-CIM tumors, respectively (|r_pb_| ≥ 0.6; p_adj_ ≤ 0.05; **Supplemental Table 4**). The top 200 differentially induced genes associated with each trajectory are displayed in **Figure 2C**. Among the top genes associated with and highly induced within Fun-CIM tumors were immune cell-activating and inflammatory signaling genes such as *IL1R1*, *TLR4*, *NLRP1*, *CD40*, *CD28*, *STAT5B*, *IRF4*, and *CCR2*. In contrast, the top genes associated with, and highly induced in Dys-CIM tumors included cytosolic and mitochondrial proteostatic and metabolic stress-adaptation genes such as *CLPP*, *PHB1*, *PHB2*, *PSMA7*, *PSMC3*, *PRDX5*, and *ATP5IF1*. To define the transcriptional programs underlying Fun-CIM and Dys-CIM tumors, we performed overrepresentation analysis of the differentially induced genes associated with each trajectory mapping to Gene Ontology (GO) Biological Process terms (**Supplemental Table S5**). Chemotherapy-induced genes in the Fun-CIM trajectory were consistently enriched for immunostimulatory and immune-activation programs with significant enrichment of adaptive immune response, including lymphocyte activation and differentiation, leukocyte proliferation and migration, cytokine production, and immune receptor signaling (**Figure 2D**). In contrast, the Dys-CIM trajectory was enriched for proteostasis and proteome maintenance programs alongside proliferative and metabolic reprogramming (**Figure 2E**). Notably, genes significantly associated with the Dys-CIM trajectory did not map to adaptive immune or antitumoral immune processes in overrepresentation analyses.

Consistent with the transcriptional programs identified above, gene set analyses performed across the 19 curated CIM-related pathways used within the CIMIC pipeline showed significant coordinated induction of selected CIM pathways related to adaptive immune response, antigen processing and presentation, T- and B-cell activation, leukocyte chemotaxis, and inflammatory signaling pathways in Fun-CIM tumors and coordinated induction of mitochondrial stress pathways in Dys-CIM tumors (**Figure 2F; Supplemental Table 6**).

### Fun-CIM tumors exhibit enhanced induction of inflammatory regulated cell death programming and CIM sensors, mediators, and downstream outputs as compared to Dys-CIM tumors

Notably among selected CIM pathways, Fun-CIM tumors demonstrated markedly higher induction of inflammatory cell apoptotic processes program (**Figure 2G**), with significantly higher induction of pyroptotic, ferroptotic, PANoptotic, and necroptotic transcriptomic programming (**Figure 2H-2K**), while there was no significant difference in the induction of apoptotic signaling (**Figure 2L**). Downstream analyses examining the induction of genes comprising programmatic signatures revealed induction of key genes related to immunogenic regulated cell death programs, including *AIM2*, *CASP1, CASP4, CASP5, GSDME, NLRP3*, *NLRC4*, and *IL18* for pyroptosis (**Supplemental Figure S1**), *ACSL4*, *PTGS2*, and *SAT1* for ferroptosis (**Supplemental Figure S2**), *AIM2*, *CASP8*, *ZBP1*, *FADD*, *RIPK3*, and *IRF1* for PANoptosis (**Supplemental Figure S3**), and *MLKL*, *RIPK3,* and *ZBP1* for necroptosis (**Supplemental Figure S4**). The induction of genes used to evaluate apoptosis is shown in **Supplemental Figure S5**.

Interestingly, Fun-CIM tumors demonstrated greater induction of *EIF2AK3* (PERK) in comparison to Dys-CIM tumors; however, they demonstrated no significant difference in the induction of *EIF2A*, and a significant decrease in the induction of *ATF4* (**Supplemental Figure S6A-S6C**). Paradoxically, Dys-CIM tumors demonstrated significantly greater induction of canonical ICD DAMPs, including *CALR*, *HMGB1*, and *PDIA3*, but no difference was observed in *ANXA1* induction (**Supplemental Figure S6E-S6H**). Alongside greater induction of the ER stress sensor *EIF2AK3*, Fun-CIM tumors demonstrated significantly greater induction of other CIM sensors, including sensors of cytosolic DNA (*IFI16*, *TLR9*, and *TLR3*, alongside previously reported *AIM2* and *ZBP1*) and RNA (*TLR3* and *TLR7*), extracellular ATP (*P2RX7*), DAMPs (*FPR1* and *TLR4*), type I IFNs (*IFNAR1* and *IFNAR2*), and IFN-γ (*IFNGR1*) (**Supplemental Figure S7**). Translators of sensing signals, such as *STING1,* were also significantly induced in Fun-CIM (**Supplemental Figure S7**). In addition, Fun-CIM tumors demonstrated greater induction of inflammatory response master regulators (*IFNG* and *NFKB1*) and antigen presentation machinery, reflected by increased average induction of MHC class I genes. Furthermore, Fun-CIM tumors demonstrated a significant coordinated induction of multiple immune checkpoint and regulatory molecules, including *PDCD1*, *CD274* (PD-L1), *BTLA*, *LAG3*, *CTLA4*, *HAVCR2* (TIM-3), and *TIGIT* as compared with Dys-CIM tumors (**Supplemental Figure S8**).

Collectively, our findings on the transcriptomic level define Fun-CIM as a chemotherapy-induced immune-activating trajectory characterized by induction of CIM-sensing machinery and compensatory checkpoint upregulation in the context of immune priming and define Dys-CIM as a chemotherapy-induced trajectory characterized by reduction of CIM-sensing machinery and heightened tumor-intrinsic stress-adaptation, proliferative capacity, and metabolic plasticity.

### CIMIC-identified CIM trajectories are associated with clinical and immunological characteristics and modulations

We next evaluated whether CIM trajectories were associated with baseline clinical and molecular characteristics. Dys-CIM tumors were significantly enriched for features of aggressive disease biology (**Table 4**). Specifically, the TNBC subtype was overrepresented in the Dys-CIM trajectory compared with the Fun-CIM trajectory (Dys-CIM: 66.7% vs. Fun-CIM: 33.3%; p = 0.0018; **Figure 3A**). Consistently, baseline PAM50 analysis of biopsy specimens demonstrated a significantly higher prevalence of basal-like tumors in the Dys-CIM group relative to Fun-CIM tumors (63.6% vs. 16%; p < 0.001; **Figure 3B**). In contrast, Fun-CIM tumors were predominantly ER-positive and enriched for Luminal A and B molecular subtypes. Moreover, post-treatment tumors exhibited trajectory-dependent molecular remodeling (**Figure 3C**). Following chemotherapy, Fun-CIM tumors tended to transition toward the less aggressive Luminal A subtype. In contrast, Dys-CIM tumors preferentially adopted or maintained a basal-like molecular phenotype. Concordant with these findings, high-grade tumors were significantly more prevalent in the Dys-CIM trajectory compared to Fun-CIM tumors (**Figure 3D**).

**Figure 3.**
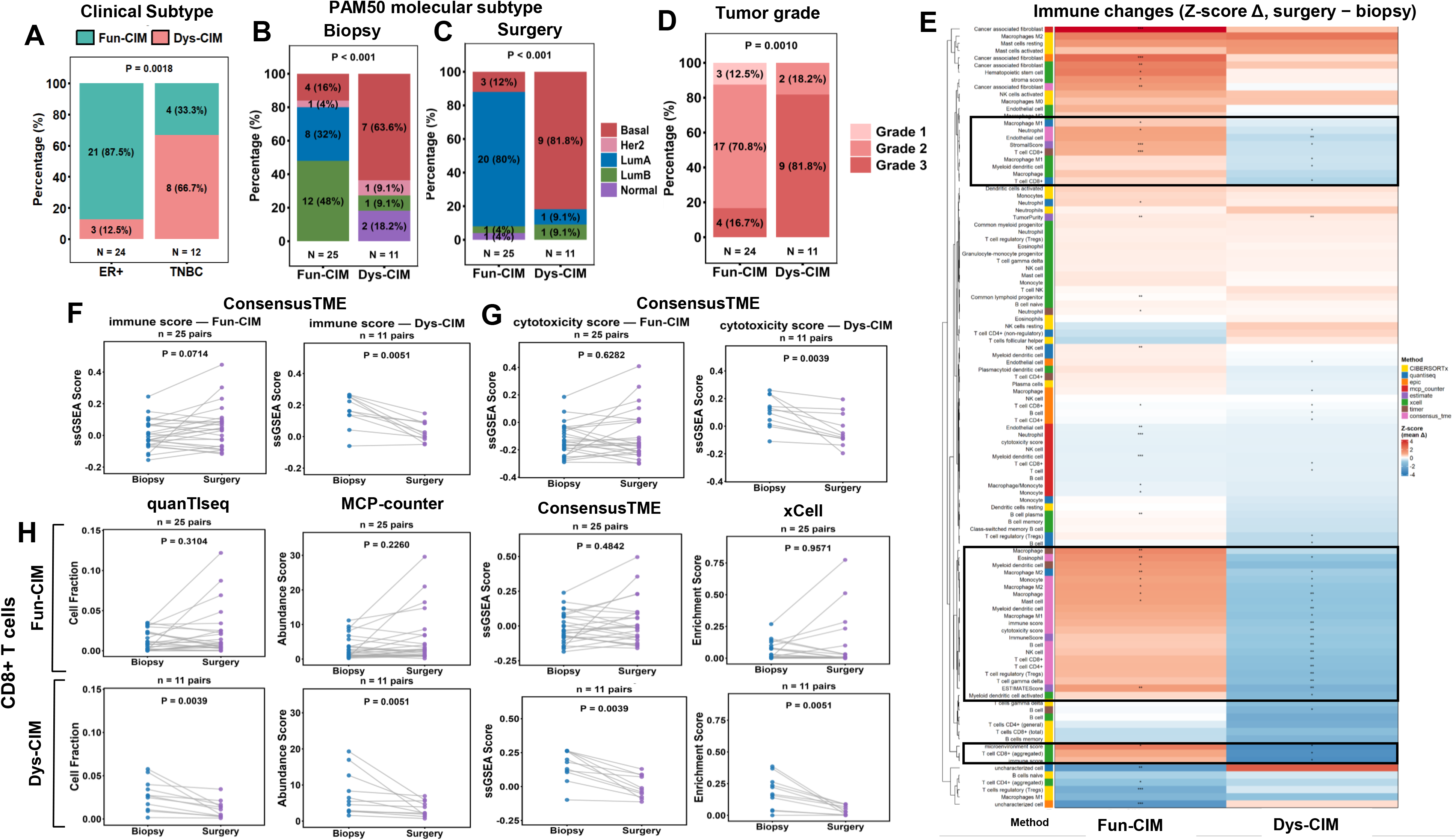
CIM trajectories associate with clinical characteristics and immune remodeling. **(A)** Baseline Fun-CIM and Dys-CIM tumor distribution across ER⁺ and TNBC subtypes, with significant overrepresentation of TNBC in Dys-CIM and ER⁺ tumors in Fun-CIM (*p* = 0.0018). **(B)** Baseline PAM50 molecular subtypes stratified by CIM trajectory showing overrepresentation of basal-like tumors in the Dys-CIM group. **(C)** PAM50 molecular subtype at surgery by CIM trajectory, with Dys-CIM tumors significantly enriched for the basal-like subtype and Fun-CIM tumors enriched for Luminal A molecular subtype. **(D)** Baseline tumor grade distribution by CIM trajectory. Dys-CIM tumors show enrichment for high-grade (Grade 3) tumors compared to Fun-CIM (*p* = 0.0010). **(E)** Heatmap of paired biopsy-to-surgery immune changes (Z-score Δ, surgery − biopsy) illustrate distinct CIM-associated immune remodeling using eight deconvolution methods. Fun-CIM tumors show induced antitumor immune infiltration, whereas Dys-CIM tumors exhibit coordinated immune contraction. Each row represents an immune cell type or immune score, and columns represent the mean cluster Z-score delta (surgery − biopsy) grouped by CIM trajectory. Paired analysis of immune **(F)** and cytotoxicity scores **(G)** by ConsensusTME shows a significant post-therapy reduction in Dys-CIM tumors. **(H)** Paired biopsy-to-surgery comparisons of CD8⁺ T-cell and abundance in Fun-CIM (N = 25, top row) and Dys-CIM tumors (N= 11, bottom row) using quanTIseq MCP-counter, ConsensusTME and xCell. Each dot represents an individual patient, with lines connecting paired samples. Significant decrease in CD8⁺ T-cell fractions and enrichment scores were detected in Dys-CIM tumors.

To investigate differences in chemotherapy-induced immune remodeling, we applied combinatorial immune deconvolution analyses integrating eight complementary deconvolution methods (**Supplemental Table 7**). The heatmap in **Figure 3E** displays divergent immune remodeling patterns across CIM trajectories, with Fun-CIM tumors exhibiting relative preservation or induction of antitumor immune infiltration, whereas Dys-CIM tumors exhibited contraction of cytotoxic immune compartments. We next focused on significant and consistent chemotherapy-induced changes in composite immune metrics using paired biopsy-to-surgery analyses. Notably, Dys-CIM tumors exhibited a significant overall reduction in immune score, reflecting estimated tumor-infiltrating lymphocyte (TIL) abundances, as detected by ConsensusTME (*p* = 0.0051) and other deconvolution methods (**Figure 3F**; **Supplemental Table 7**). Concordantly, Dys-CIM tumors exhibited reduced cytotoxic activity following chemotherapy, as detected by a decrease in the ConsensusTME cytotoxicity score (*p* = 0.0039, **Figure 3G**) and significant reductions in CD8⁺ T-cell (**Figure 3H**) and NK cell abundances (**Supplemental Figure S9**) across multiple immune deconvolution methods.

### CIM trajectory-specific genes are significantly associated with outcome in breast cancer patients treated with chemotherapy

To assess the clinical relevance of CIM trajectory-associated genes, we examined whether biopsy-to-surgery changes in the expression of genes associated with the Fun-CIM (*n* = 542) and Dys-CIM (*n* = 269) trajectories were associated with RFS in the NKI cohort. In univariable Cox analyses, 53 of 542 Fun-CIM genes (9.8%) were classified as beneficial (HR < 1 and p < 0.05), whereas only three were classified as detrimental (HR > 1 and p < 0.05). Conversely, 223 of 269 Dys-CIM genes (82.9%) were classified as detrimental, and none were classified as beneficial (**Figure 4A**; **Supplementary Table 8**)

**Figure 4.**
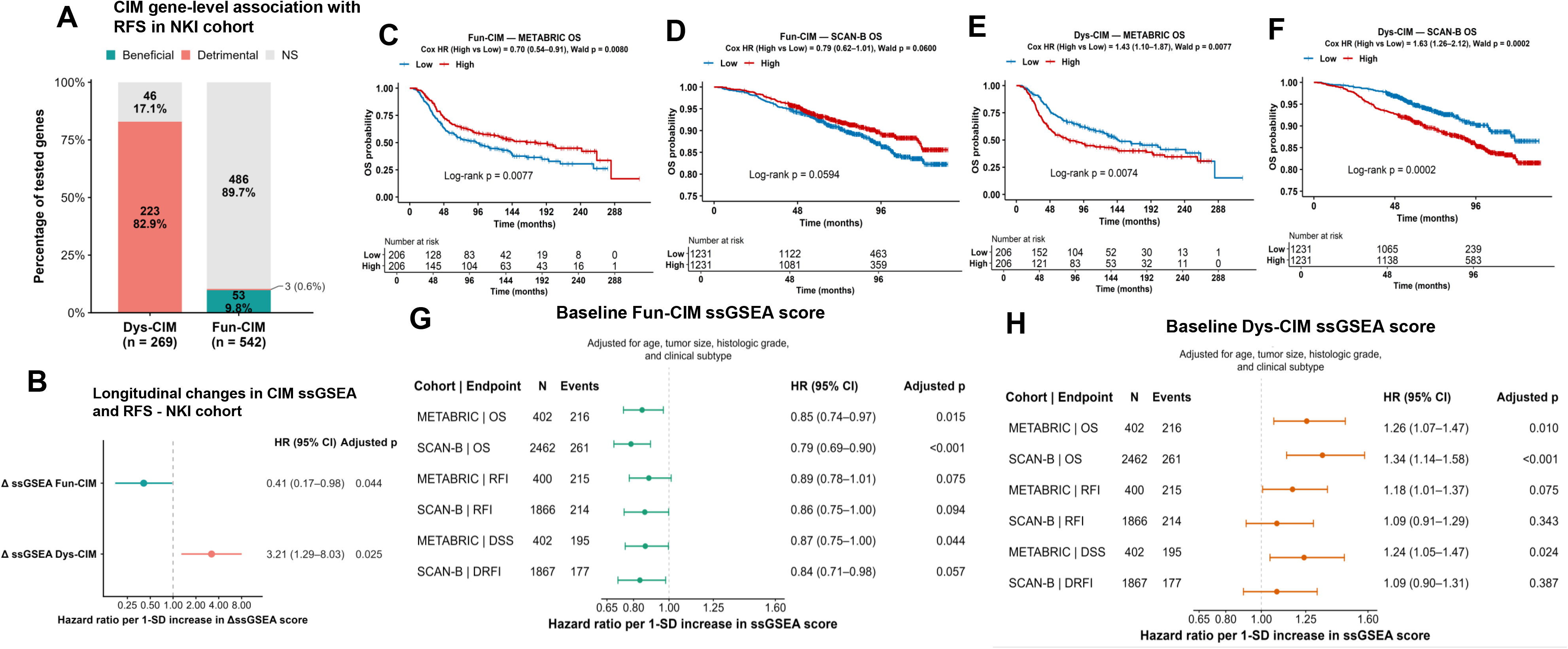
Longitudinal and baseline clinical relevance of Fun-CIM and Dys-CIM transcriptional programs. **(A)** Directional distribution of gene-level associations with recurrence-free survival (RFS) in the NKI cohort. Trajectory-specific gene sets composed of genes whose chemotherapy-induced delta expression was positively correlated with Dys-CIM (n = 269) and Fun-CIM (n = 542) trajectories (Point biserial correlation |r_pb_| *≥* 0.60, FDR adjusted p < 0.05) were used. For each gene, change was calculated from Δlog₂(TPM+1) expression as surgery minus biopsy and standardized within gene. Each gene was evaluated separately using a univariable Cox proportional-hazards model; Genes with (nominal p < 0.05) and HR < 1 were classified as beneficial, whereas genes with (nominal p < 0.05) and HR > 1 were classified as detrimental; all others were classified as not significant (NS) with HRs represent the association per 1-SD greater biopsy-to-surgery expression change. **(B)** Associations between longitudinal changes in Fun-CIM and Dys-CIM ssGSEA scores and RFS in NKI. Hazard ratios represent the association per one-standard-deviation (1-SD) increase in the ΔssGSEA change. **(C–F)** Kaplan–Meier analyses stratified by cohort- and signature-specific median baseline ssGSEA scores. High Dys-CIM was associated with worse overall survival (OS) in METABRIC **(C)** and SCAN-B **(D),** whereas High Fun-CIM was associated with improved OS in METABRIC **(E)** and showed a favorable trend in SCAN-B **(F).** Cox HRs with 95% CIs and log-rank (p) values are shown. **(G,H)** Multivariable Cox analyses of continuous baseline Dys-CIM **(G)** and Fun-CIM **(H)** ssGSEA scores across OS, disease-specific survival (DSS), recurrence-free interval (RFI), and distant recurrence-free interval (DRFI) in chemotherapy-treated METABRIC and SCAN-B patients. Models were adjusted for age, tumor size, histologic grade, and clinical subtype. HRs are reported per 1-SD increase in ssGSEA score; horizontal lines indicate 95% CIs, and the dashed line indicates HR = 1. Higher Dys-CIM was independently associated with worse OS in both cohorts and worse DSS in METABRIC, whereas higher Fun-CIM was associated with improved OS in both cohorts and improved DSS in METABRIC. HR, hazard ratio; CI, confidence interval; ssGSEA, single-sample gene set enrichment analysis.

We next calculated sample-level Fun-CIM and Dys-CIM ssGSEA scores and evaluated whether their biopsy-to-surgery changes (ΔssGSEA) were associated with RFS in the same cohort. In an exploratory analysis, a greater increase in the Fun-CIM ssGSEA score was associated with improved RFS (HR per 1-SD increase = 0.41, 95% CI 0.17-0.98, adjusted p = 0.044), whereas a greater increase in the Dys-CIM score was associated with worse RFS (HR = 3.21, 95% CI 1.29-8.03, adjusted p = 0.025; **Figure 4B**). Thus, both gene-level and signature-level analyses showed directionally opposing associations of the Fun-CIM and Dys-CIM transcriptional programs with clinical outcomes.

We then evaluated the prognostic relevance of baseline Fun-CIM and Dys-CIM ssGSEA scores among chemotherapy-treated patients in METABRIC (up to *n* = 412) and SCAN-B (up to *n* = 2,462) cohorts. In univariable Cox analyses using cohort- and signature-specific median stratification, high Fun-CIM scores were associated with improved OS in METABRIC (HR = 0.70, 95% CI 0.54-0.91, p = 0.008) and showed a favorable trend in SCAN-B (HR = 0.79, 95% CI 0.62-1.01, p = 0.06; **Figure 4E-4F**). Conversely, high Dys-CIM scores were associated with worse OS in METABRIC (HR = 1.43, 95% CI 1.10-1.87, p = 0.0077) and SCAN-B (HR = 1.63, 95% CI 1.26-2.12, p < 0.001; **Figure 4C-4D**).

Consistent directional associations were also observed across other endpoints. High Fun-CIM signature scores were associated with improved DSS in METABRIC and improved RFI and DRFI in SCAN-B, whereas high Dys-CIM signature scores were associated with worse DSS and RFI in METABRIC and worse RFI and DRFI in SCAN-B (**Supplementary Table 9**).

In multivariable Cox models adjusted for age, tumor size, histologic grade, and clinical subtype, higher continuous Fun-CIM scores were associated with improved OS in METABRIC (HR per 1-SD increase = 0.85, 95% CI 0.74-0.97, adjusted p = 0.015) and SCAN-B (HR = 0.79, 95% CI 0.69-0.90, adjusted p < 0.001), as well as improved DSS in METABRIC (HR = 0.87, 95% CI 0.75-1.00, adjusted p = 0.044; **Figure 4H; Supplementary Table 10**). Conversely, higher continuous Dys-CIM ssGSEA scores were associated with worse OS in METABRIC (HR = 1.26, 95% CI 1.07-1.47, BH-adjusted p = 0.010) and SCAN-B (HR = 1.34, 95% CI 1.14-1.58, adjusted p < 0.001), as well as worse DSS in METABRIC (HR = 1.24, 95% CI 1.05-1.47, adjusted p = 0.024; **Figure 4G**; **Supplementary Table 11**). Multivariable models using cohort-specific median stratification yielded concordant results and showed significant favorable RFI and DRFI associations for high Fun-CIM in SCAN-B (**Supplementary Tables 12** and **13**).

### Validation of CIM trajectories derived from CIMIC in the NEO dataset

To assess the applicability and generalizability of CIMIC, we applied CIMIC to paired pre- and post-treatment data from the NEO study. An optimal solution was identified at k = 2 and determined to be distinct and stable based on our composite clustering metrics framework (**Figure 5A**). Heatmap visualization of ΔGE across all protein-coding genes revealed distinct induction patterns in response to chemotherapy between Fun-CIM and Dys-CIM tumors, characterized by broad induction in Fun-CIM tumors and comparatively limited transcriptional remodeling in Dys-CIM tumors (**Figure 5B**), a phenotype qualitatively distinct from the active proteostatic stress-adaptive remodeling observed in NKI/SMC Dys-CIM tumors. Accordingly, point-biserial correlation derived from limma’s moderated *t*-statistics between Fun-CIM and Dys-CIM tumors identified 3,436 differentially induced genes, with 3,427 genes associated with Fun-CIM and 9 associated with Dys-CIM tumors (|r_pb_| ≥ 0.6, p_adj_ ≤ 0.05; **Supplemental Table 14**). Notably, the top genes differentially induced in Fun-CIM tumors included immune signaling genes such as *STING1, RELA, IL17RA, LITAF, LTBR, MAPK14,* and *TNFR1* (**Figure 5C**), similar to Fun-CIM-associated genes identified in the NKI/SMC cohort. In overrepresentation analysis, genes significantly differentially induced in Fun-CIM tumors identified inflammatory cellular stress response and immune activation pathways, including interferon-beta production, inflammasome-mediated signaling, activation of innate immune responses, and viral mimicry-like signaling (**Figure 5D**). In contrast, genes significantly induced in Dys-CIM tumors were predominantly from the small nucleolar RNA (snoRNA) gene family, which is critical for pseudouridylation of ribosomal RNA, a key process involved in ribosomal RNA modification and ribosome biogenesis (**Figure 5C**). No meaningful results were produced using these genes for overrepresentation analysis (**Supplemental Table 15**).

**Figure 5.**
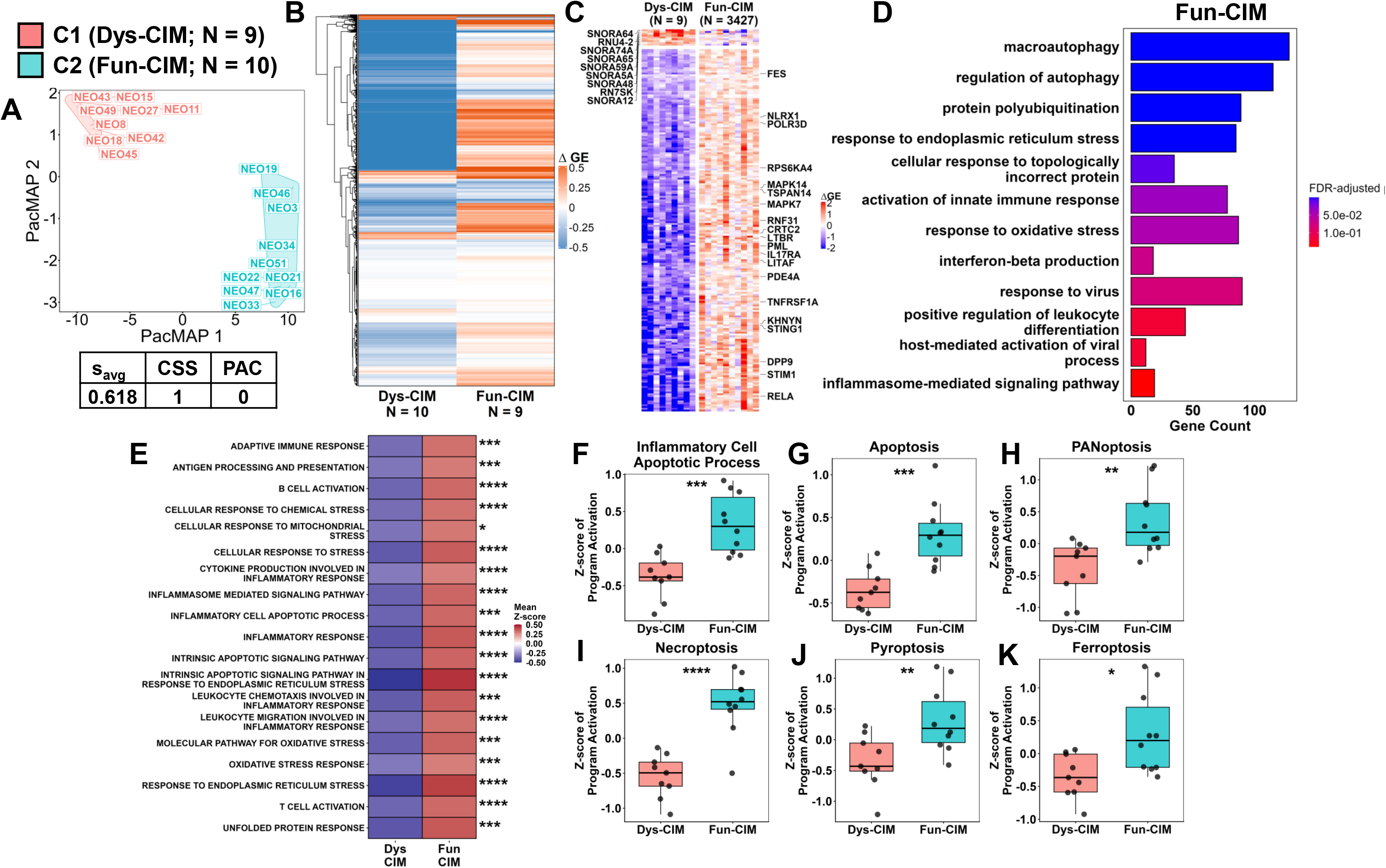
CIMIC identifies divergent transcriptional CIM trajectories in the NEO cohort. **(A)** PaCMAP projection of ΔGE demonstrating segregation of breast cancer patients into two distinct CIM trajectories: C1 (Dys-CIM; N = 9) and C2 (Fun-CIM; N = 10). Average silhouette score from ΔGE and PaCMAP reduced space (s_avg_), cluster consensus score (CSS), and proportion of ambiguous clustering (PAC) metrics are indicated. **(B)** Heatmap of ΔGE across all protein-coding genes, revealing distinct and opposing transcriptional responses to chemotherapy between Fun-CIM and Dys-CIM trajectories. **(C)** Heatmap of the top 200 CIM-trajectory associated genes (|r_pb_| ≥ 0.6; FDR- adjusted p ≤ 0.05) for Fun-CIM and Dys-CIM trajectories. **(D)** Top enriched Gene Ontology (GO) biological processes among genes preferentially induced in Fun-CIM tumors following chemotherapy (FDR-adjusted p < 0.15). No meaningful pathways rose to the level of significance in Dys-CIM tumors (**Supplemental Table 15**). **(E)** Heatmap depicts the relative differential induction (Z-score of average ΔGE values) of selected chemoimmunomodulatory pathways. **(F-K)** Boxplots comparing the relative differential induction (Z-score of average ΔGE values) of regulated cell-death programs between Fun-CIM and Dys-CIM tumors, including **(F) I**nflammatory cell apoptotic process, **(G)** Apoptosis, **(H)** PANoptosis **(I)**, Necroptosis **(J)**, and **(K)** Ferroptosis. Fun-CIM tumors exhibited significantly greater induction of all evaluated cell-death programs compared with Dys-CIM tumors. FDR-adjusted p values are indicated as follows: * ≤ 0.05; ** ≤ 0.01; *** ≤ 0.005; **** ≤ 0.001; ns, not significant.

Consistent with the broad transcriptional induction observed in Fun-CIM tumors and comparatively limited remodeling in Dys-CIM tumors, gene set analyses of CIM-related pathways showed broad induction of CIM-related programs, including significant induction of adaptive immune response, antigen processing and presentation, T- and B-cell activation, leukocyte chemotaxis, and inflammatory signaling pathways in Fun-CIM tumors (**Figure 5E; Supplemental Table 15**). Similarly, Fun-CIM tumors demonstrated higher induction of inflammatory cell apoptotic processes (**Figure 5F**), with significantly higher induction of apoptotic, PANoptotic, necroptotic, and pyroptotic transcriptomic programming (**Figure 5G-5K**).

An analysis of CIM sensors and signal translators revealed generally higher induction in Fun-CIM tumors as compared to Dys-CIM tumors, with significantly higher induction of *STING1*, *TLR9*, and *IFNAR1* observed (**Supplemental Figure S10**). While Fun-CIM tumors did have higher induction of *NFKB1* and *VSIR*, no significant differences were observed in the induction of other immune checkpoints and MHC class I genes (**Supplemental Figure S11**). Similarly, there were no significant differences in the distribution of PAM50 subtypes across Fun-CIM and Dys-CIM tumors (**Supplemental Figure S12A-S12B**). Despite differences in transcriptomic reprogramming, immune deconvolution analyses did not reveal substantial differences in inferred immune cell composition between Fun-CIM and Dys-CIM tumors (**Supplemental Figure S12C-S12E**). Nonetheless, these findings demonstrate the ability of CIMIC to identify divergent Fun-CIM and Dys-CIM trajectories in an independent dataset. Notably, the Dys-CIM trajectory within the NEO cohort demonstrated limited transcriptional remodeling in response to chemotherapy, suggesting that Dys-CIM may not represent a single invariant transcriptional state but rather a spectrum of context-dependent chemotherapy-induced responses, ranging from active tumor-intrinsic stress-adaptation to blunted immunomodulatory responses, or combinations thereof.

### Orthogonal in vitro corroboration of tumor-intrinsic CIM programs using triple-negative breast cancer cell lines

As an orthogonal validation of our findings in patients and to assess the extent to which divergent CIM trajectories are driven by tumor-intrinsic biology, we applied a modified version of CIMIC using only tumor cell-relevant programs to paired pre- and post-treatment RNA-seq data from 9 TNBC cell lines, with post-treatment samples subjected to CTG-derived IC30 concentrations of epirubicin to standardize sublethal cytotoxic perturbation (**Figure 6A-6B**). As in patients, an optimal, stable, and distinct solution was identified at k = 2 (**Figure 6C**), with distinct global induction patterns observed across clusters (**Figure 6D**). Point-biserial correlation derived from limma’s moderated *t*-statistics between Fun-CIM and Dys-CIM cell lines identified 863 differentially induced genes, with 408 genes associated with Fun-CIM and 455 associated with Dys-CIM cell lines, respectively (|r_pb_| ≥ 0.6; p_adj_ ≤ 0.05; **Figure 6E; Supplemental Table 17**). As in patient tumors, Fun-CIM cell lines demonstrated significantly increased induction of immune signaling molecules including *STING1, STAT2,* and *IFI35,* as well as significantly increased induction of early stress-response genes, including *FOS*, *EGR1*, and *FOSB*. In contrast, Dys-CIM cell lines had induction of proteostatic adaptation genes, including several members of the ubiquitin-proteasome system (*USP7, UBE3C, UBQLN1, PSMD14, SMURF2, FAF1*), mitochondrial maintenance and metabolism genes (*ATP5F1A, CHCHD3, TAMM41, VDAC, TRAP1*), and epithelial-mesenchymal transition (*ZEB1*, *ZEB2, WWTR1*) genes. Although differences in the induced gene programs were less distinct between Fun-CIM and Dys-CIM cell lines, overrepresentation analysis of genes associated with the Fun-CIM cell lines mapped to inflammatory cellular stress response programs, such as response to interleukin-1, and viral mimicry-like signaling as in patients (**Supplemental Figure S13A**). In contrast, genes significantly differentially induced in Dys-CIM cell lines mapped to proteome maintenance and proteostatic stress response pathways and immunosuppressive signaling programs, such as response to transforming growth factor-beta (**Supplemental Figure S13B; Supplemental Table 18**). Gene set analyses of CIM-related pathways revealed that Fun-CIM tumors had significantly greater induction in programs like antigen presentation and processing and inflammatory cell apoptosis as compared to Dys-CIM, although this was not statistically significant (**Figure 6F**). In contrast, Dys-CIM cell lines had significantly greater induction of the cellular response to mitochondrial stress program, corroborating our findings in patients.

**Figure 6.**
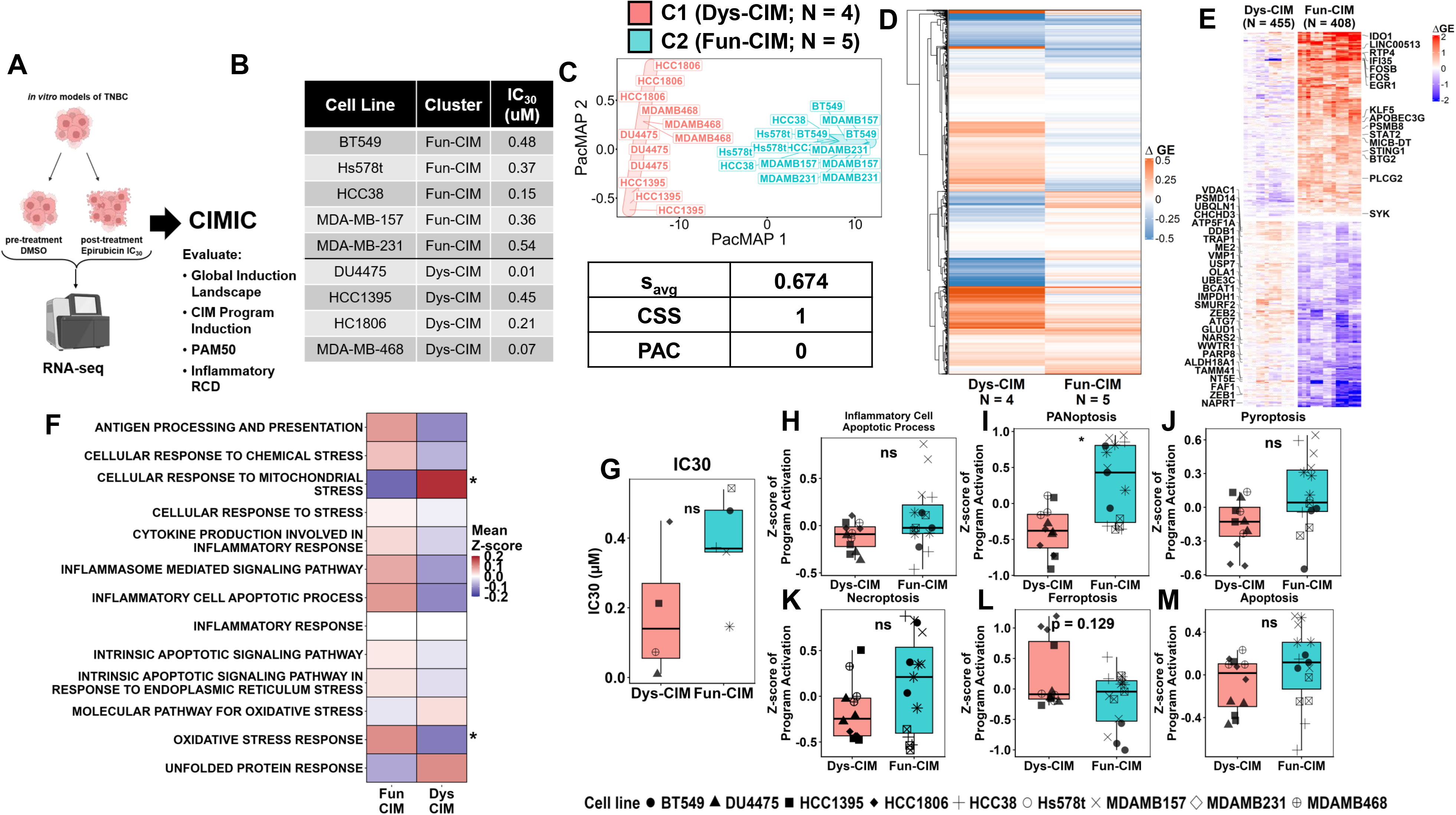
CIMIC captures divergent immunomodulatory programming induced by chemotherapy in TNBC cell lines. **(A)** Schematic workflow of the experimental design: TNBC cell lines were treated with either DMSO (pre-treatment) or epirubicin (post-treatment) at their respective IC_30_ concentrations as determined by CellTiterGlo for 48 hours, followed by RNA-sequencing and classification of CIM trajectories using CIMIC followed by assessment global induction landscapes, program induction, and regulated cell death (RCD) pathways between identified CIM trajectories. Samples were collected and analyzed in biological triplicates (see Methods section for detail). **(B)** Summary table of triple-negative breast cancer (TNBC) cell lines utilized for *in vitro* validation, included their IC_30_ values for epirubicin used in downstream experiments. **(C)** PaCMAP projection of ΔGE demonstrating segregation of TNBC cell line samples into two distinct CIM trajectories: C1 (Dys-CIM; N = 4 cell lines; N = 12) and C2 (Fun-CIM; N = 5 cell lines; N = 15). Average silhouette score from ΔGE and PaCMAP reduced space (s_avg_), cluster consensus score (CSS), and proportion of ambiguous clustering (PAC) metrics are indicated. **(D)** Heatmap of ΔGE across all protein-coding genes, revealing distinct and opposing transcriptional responses to chemotherapy between Fun-CIM and Dys-CIM trajectories within the cell line dataset. **(E)** Heatmap of the top 200 CIM-trajectory associated genes (|r_pb_| ≥ 0.6; FDR-adjusted p ≤ 0.05) for Fun-CIM and Dys-CIM trajectories. **(F)** Heatmap depicts the relative differential induction (Z-score of average ΔGE values) of selected chemoimmunomodulatory pathways used for clustering in Fun-CIM and Dys-CIM cell lines. Box plot depicts comparison of epirubicin IC_30_ sensitivity between Dys-CIM and Fun-CIM cell lines **(G)** and the relative differential induction (Z-score of average ΔGE values) of regulated cell death programs in Fun-CIM (blue) and Dys-CIM (red) cell lines **(I-L)**. Fun-CIM tumors demonstrated a greater, non-significant induction of inflammatory cell death programming **(H)** and significantly greater induction of PANoptotic programming **(I)**. There were no significant differences in pyroptotic, necroptotic, ferroptotic, and apoptotic programming. Statistically significant differences for FDR-adjusted p values in **(F)** and **(I-L)** are indicated as follows: * ≤ 0.05; ** ≤ 0.01; *** ≤ 0.005; **** ≤ 0.001.

Interestingly, although there were no significant differences in epirubicin sensitivity between Fun-CIM and Dys-CIM tumors (**Figure 6G**), Fun-CIM cell lines demonstrated a greater, although non-significant, induction of inflammatory cell death programming (**Figure 6H**), including significantly greater induction of PANoptotic programming (**Figure 6I**). No significant differences were observed in pyroptotic, necroptotic, ferroptotic, and apoptotic programming. Similar to patients, Fun-CIM cell lines generally had higher induction of immune checkpoint molecules, including significant induction of *PDCD1*, *CTLA4*, and other B7 family members (**Supplemental Figure S14**). Given our findings in differential gene induction analyses and the observed induction of viral mimicry-like programs across Fun-CIM patient tumors and cell lines (**Figure 2C**, **Figure 5D**, **Supplemental Figure S15**), we further investigated differences in the induction of viral mimicry programming between Fun-CIM and Dys-CIM tumors and cell lines, deriving a 47-gene signature comprising sensors, signal transducers, and functional outputs related to viral mimicry. Applying gene set activity and coordination analyses to this 47-gene signature revealed significantly greater induction of viral mimicry programming in Fun-CIM tumors and cell lines as compared to Dys-CIM tumors and cell lines (**Supplemental Figure S15A-S15F**), with notably higher induction of key cytosolic nucleic acid sensors such as *CGAS*, *STING1*, *AIM2*, *IFIH1*, *DHX58*, and *RIGI* (**Supplemental Figure S15G-S15L**) in Fun-CIM cell lines.

### Conserved transcriptional programs distinguish Fun-CIM and Dys-CIM across clinical and experimental systems

To determine whether the transcriptional remodeling associated with CIM trajectories was conserved across clinical and experimental systems, we applied Stouffer’s method to the results from our differential gene induction analyses across datasets, using a relaxed threshold of |r_pb_| ≥ 0.3 to maximize representation of directionally consistent genes, and requiring directional concordance across NKI/SMC tumors and TNBC cell lines for statistical significance. Using this approach, 1,079 genes demonstrated conserved induction across Fun-CIM tumors and cell lines, including many immunostimulatory and inflammatory genes, with overrepresentation analysis revealing conserved enrichment of immune activation and inflammatory signaling programs (**Figure 7A-7B**). In contrast, 402 genes demonstrated conserved induction across Dys-CIM tumors and cell lines, including many proteostatic and mitochondrial stress-adaptation and metabolic reprogramming genes, with overrepresentation analysis revealing conserved enrichment of proteome maintenance and proteostatic stress-adaptation programs (**Figure 7C-7D**). Similarly, conservation analysis revealed directionally concordant induction for key tumor-intrinsic immunostimulatory CIM programs including antigen presentation and processing, cytokine production, and inflammatory cell death programs (pyroptosis, PANoptosis, and necroptosis) across Fun-CIM samples. Conversely, Dys-CIM samples demonstrated directionally concordant induction of cellular stress response programs, with significant cross-dataset consensus observed specifically for cellular response to mitochondrial stress (**Figure 7E-7F**).

**Figure 7.**
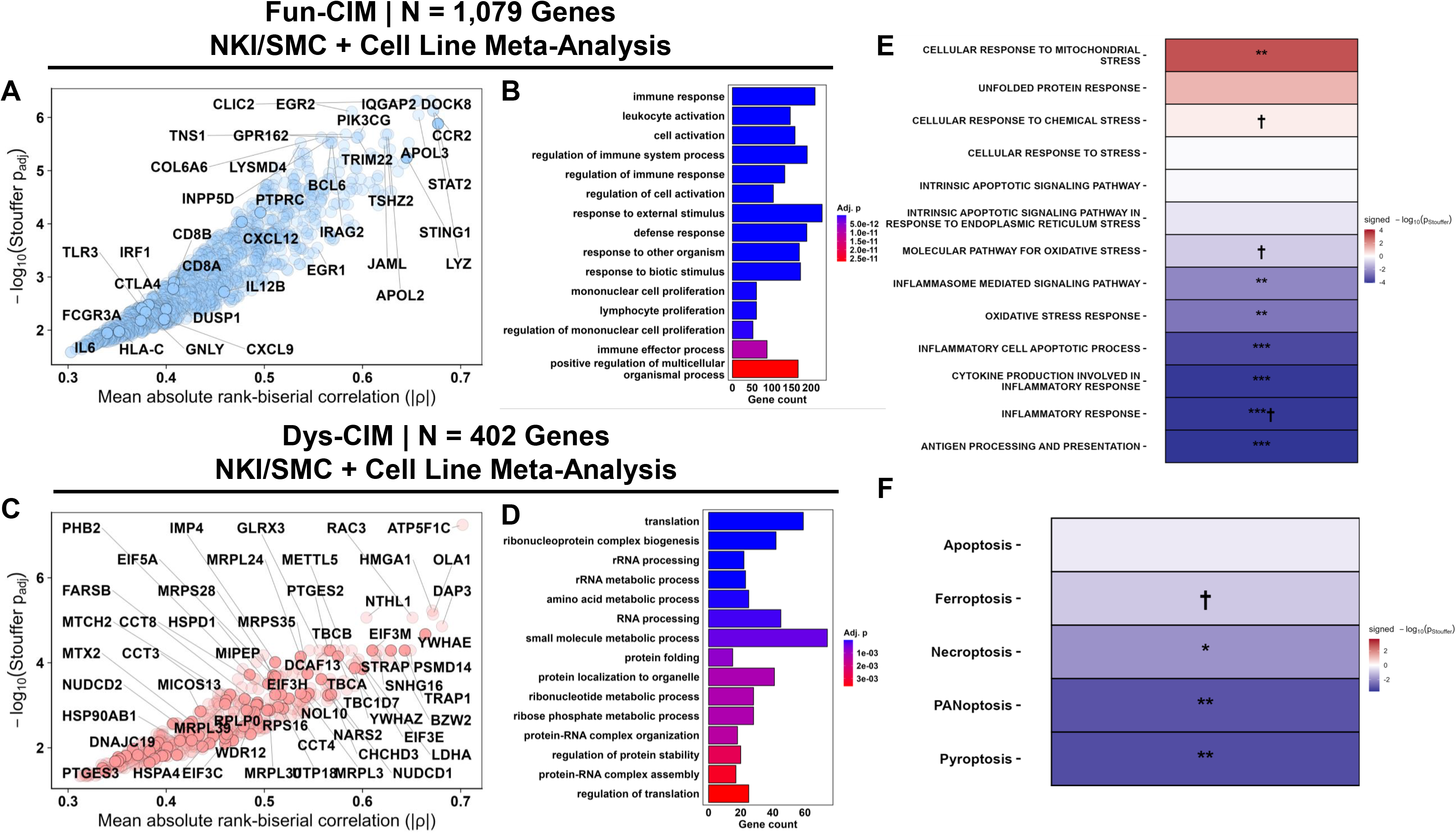
Conserved Fun-CIM and Dys-CIM transcriptional programs across patient tumors and TNBC cell lines. **(A, C)** Cross-dataset analysis of directionally concordant genes using Stouffer’s method across NKI/SMC tumors and TNBC cell lines identified 1,079 Fun-CIM (**A**) and 402 Dys-CIM (**C**) conserved genes. **(B, D)** Gene Ontology Biological Process overrepresentation analysis of conserved genes associated with **(B)** Fun-CIM and **(D)** Dys-CIM trajectories. **(E–F)** Cross-dataset analysis of CIM-related (**E**) and regulated cell-death (**F**) pathways demonstrated conserved induction of immunostimulatory and inflammatory cell-death programs in Fun-CIM, whereas Dys-CIM showed greater conservation of cellular stress responses. Heatmap colors represent signed −log10(Stouffer p *value*s).

Although the Dys-CIM trajectory in the NEO cohort exhibited a more transcriptionally limited phenotype as compared to NKI/SMC and TNBC cell lines, we performed a sensitivity conservation analysis integrating all three datasets. The Fun-CIM trajectory demonstrated substantial cross-dataset conservation, with 551 directionally concordant genes conserved across all three datasets and an enrichment of immune activation and inflammatory signaling programs (**Supplemental Figure S16A-16B**). In contrast, only 6 genes were directionally concordant and conserved across all three datasets for the Dys-CIM trajectory, consistent with the greater heterogeneity in the Dys-CIM trajectory observed across datasets. Nevertheless, overrepresentation analysis using a relaxed FDR threshold (p_adj_ < 0.15) revealed enrichment of protein maintenance programs, consistent with the conserved Dys-CIM biology observed in the NKI/SMC and TNBC cell line analysis (**Supplemental Figure S16C-16D**). At the pathway level, CIM-related programs and inflammatory cell death programs demonstrated directionally concordant patterns across all three datasets, including directionally concordant conserved induction of immunostimulatory and inflammatory cell death programs observed in the Fun-CIM trajectory as opposed to the Dys-CIM trajectory (**Supplemental Figure S16E-16F**).

To assess the clinical relevance of conserved Fun-CIM and Dys-CIM genes identified across multiple datasets, we evaluated the association between biopsy-to-surgery ΔGE of genes identified in the two- and three-dataset conservation analyses and RFS in the NKI cohort. Both biopsy-to-surgery gene-level expression changes and signature-level ssGSEA changes demonstrated significant associations with RFS (**Supplementary Figure S17**), supporting the clinical relevance of the conserved transcriptional programs associated with the Fun-CIM and Dys-CIM trajectories.

Collectively, these findings demonstrate that CIMIC identifies Fun-CIM and Dys-CIM trajectories in tumor cells under controlled chemotherapy exposure and that key components of these trajectories are conserved between patient tumors and tumor cell models. This concordance across clinical and experimental systems suggests a tumor-intrinsic dimension of CIM in which chemotherapy-induced programs within tumor cells contribute to the broader immunomodulatory trajectory of the tumor and demonstrates that the classification of Fun-CIM and Dys-CIM defines stable and biologically meaningful transcriptional trajectories.

### Baseline differences between Fun-CIM and Dys-CIM tumors and cell lines

To assess the extent to which CIM induction was determined by pre-treatment transcriptional states, we first evaluated the correlation between baseline CIM gene expression and ΔGE. These correlations were weak across patient cohorts and TNBC cell lines (**Supplemental Methods SM1**). Nonetheless, we investigated whether Fun-CIM and Dys-CIM trajectories arose from distinct pre-treatment transcriptional states by comparing baseline gene expression between the two groups across cohorts. No significant baseline differentially expressed genes were identified between Fun-CIM and Dys-CIM in the NEO cohort or TNBC cell lines (data not shown). In contrast, analysis of the combined NKI and SMC cohorts identified 1,990 genes with higher baseline expression in Fun-CIM tumors and 709 genes with higher expression in Dys-CIM tumors (**Supplemental Table 19**). Notably, baseline Dys-CIM tumors were enriched for adaptive immunity, lymphocyte activation, leukocyte-mediated immunity and cell killing, whereas baseline Fun-CIM tumors were enriched for microtubule- and cilium-related processes and intracellular transport (**Supplemental Figure S18A-S18B**).

We therefore examined whether the immune-related transcriptional enrichment observed in Dys-CIM was accompanied by differences in baseline immune composition. Dys-CIM tumors also exhibited significantly higher immune scores by xCell (p = 0.0110) and ESTIMATE (p = 0.0009) and greater CD8^+^ T-cell abundance across multiple deconvolution methods, whereas differences in NK-cell abundance were inconsistent. (**Supplemental Figure S18C-E**). Moreover, baseline expression of MHC class I expression and multiple immune checkpoints, including *PDCD1*, *CD274*, *TIGIT*, *LAG3*, *PDCD1LG2*, and *CTLA4*, was also significantly higher in Dys-CIM tumors (**Supplemental Figure S19**). Collectively, these findings indicate that Dys-CIM tumors exhibited a comparatively immune-enriched baseline state characterized by increased CD8⁺ T-cell fraction, MHC-I expression, and immune-checkpoint expression. Despite greater baseline immune enrichment, Dys-CIM tumors showed limited induction of adaptive or antitumoral immune processes following chemotherapy, whereas Fun-CIM tumors preferentially induced adaptive immunity, antigen presentation, and inflammatory signaling.

Importantly, baseline differences between Fun-CIM and Dys-CIM were not detected in the NEO cohort or TNBC cell lines, indicating that CIM trajectories may not be able to be consistently inferred from pre-treatment transcriptional features alone. Taken together, these results suggest CIM trajectories identified by CIMIC capture divergent chemotherapy-associated remodeling can arise from distinct pretreatment states, but are not consistently determined by baseline transcriptional features alone.

## 4. Discussion

CIM has become a crucial area of focus as chemotherapy is recognized as a modulator of tumor-immune interactions rather than a purely cytotoxic intervention (44,45). While CIM has numerous clinically significant implications, its effects are highly heterogeneous and context-dependent (21,23), warranting a comprehensive framework to globally resolve and classify divergent CIM trajectories. Until now, however, no such framework has been established. Here, we present CIMIC, a novel framework that addresses this gap by using transcriptional changes from pre- and post-treatment specimens to classify tumors according to their CIM trajectory. Across pre- and post-treatment data from two independent patient cohorts (NKI/SMC, N = 36; NEO, N = 19) and within an *in vitro* experimental framework where TNBC cell lines (N = 9) were subjected to standardized epirubicin perturbation, CIMIC identified two distinct, stable divergent CIM trajectories: a functional CIM trajectory, which was antitumoral and immune-activating, and a dysfunctional CIM trajectory, which was stress-adaptive and immune/blunted.

The Fun-CIM trajectory was characterized by coordinated induction of inflammatory cell death programs and CIM/innate immune sensing machinery alongside the induction of antigen presentation and adaptive immune programs, which is consistent with the productive engagement of the ICD-to-immunity cascade (3). Consistent with this immunostimulatory phenotype, Fun-CIM patient tumors and cell lines demonstrated significantly greater induction of viral mimicry programming, which is recognized as an innate immune-activating mechanism that can amplify antitumor immunity and critical bridge between ICD signaling and innate immune responses against tumors (46,47). Across patients and cell lines, Fun-CIM samples also demonstrated various compensatory upregulation of immune checkpoint receptors, including both tumor-intrinsic and tumor-extrinsic *PDCD1* and *CTLA-4* (48,49), *CD274* (PD-L1), *VSIR,* and other B7 family members, suggesting that chemotherapy drives evolution toward a tumor profile characterized by simultaneous immune activation and adaptive immune resistance that may represent a context in which tumors are more responsive to chemoimmunotherapy using immune checkpoint blockade (50–52). In the NKI/SMC cohort, immune deconvolution analyses revealed the preservation/increase of cytotoxic immune populations and activity and inferred abundance of tumor-infiltrating lymphocytes (TILs) in Fun-CIM tumors compared with Dys-CIM tumors across multiple complementary algorithms, supporting a functional link between CIM trajectory and the composition of the TME. Altogether, these findings support a model in which Fun-CIM tumors exhibit efficient coupling between the induction and sensing of chemotherapy-induced stress and downstream antitumoral immune response.

In contrast, Dys-CIM tumors preferentially induced stress-adaptive proteostatic, metabolic, and proliferative programs defined by enrichment of oxidative phosphorylation, DNA repair pathways, and regulation of ribosome structure and translational capacity and fidelity following chemotherapy. This transcriptional profile suggests that Dys-CIM tumors respond to cytotoxic perturbation by activating cell-autonomous survival machinery related to proteome maintenance and proliferative and metabolic preprograming rather than propagating chemotherapy-induced stress into immune signaling (20,53). Alongside this, the Dys-CIM trajectory was enriched for aggressive disease features, including overrepresentation of TNBC and transitioning toward basal molecular subtypes upon treatment with chemotherapy. In the NKI/SMC cohort, Dys-CIM tumors also exhibited reduced estimated TIL abundances and decreased CD8⁺ T-cell and NK-cell fractions and cytotoxic activity. Interestingly, our analyses revealed a paradoxical finding that Dys-CIM tumors showed greater induction of canonical ICD DAMPs including *CALR*, *HMGB1*, and *PDIA3*, despite limited immunostimulatory remodeling. However, Dys-CIM tumors also exhibited reduced induction or reduction of key CIM sensors for various DAMPs, suggesting a signaling bottleneck in which tumor-intrinsic stress is not translated into antitumoral immunity. Indeed, it is likely that Dys-CIM tumors may leverage the induction of these DAMPs to further drive stress-adaptation, given that several DAMPs, such as CALR and HMGB1, have established roles in proteostasis and tumor cell survival apart from their DAMP functions (54–58). This mechanistic hypothesis requires functional validation but offers a conceptually compelling explanation for why DAMP induction does not guarantee an immunogenic outcome. Importantly, characterization of Dys-CIM tumors within the NEO cohort suggests that Dys-CIM may encompass heterogeneous transcriptional manifestations rather than a single molecular state. Whereas Dys-CIM in NKI/SMC was characterized by active proteostatic and metabolic stress-adaptation, NEO Dys-CIM exhibited comparatively limited transcriptional remodeling, suggesting that failure to mount productive immunostimulatory remodeling may arise through distinct mechanisms or varying degrees of tumor-intrinsic stress-adaptation. The observed heterogeneity in Dys-CIM warrants further investigation in larger, independent cohorts to determine whether distinct molecular manifestations of Dys-CIM represent reproducible and therapeutically relevant states. Nonetheless, these findings altogether support a model in which Dys-CIM tumors preferentially engage tumor-intrinsic stress-adaptation programs upon treatment with chemotherapy while failing to propagate chemotherapy-induced stress into productive immune signaling.

An important question arising from these findings is whether divergent CIM trajectories are driven solely by interactions within the immune components of the TME or whether they also reflect tumor-intrinsic responses to chemotherapy. To address this, we applied CIMIC to TNBC cell lines under controlled epirubicin perturbation. In TNBC cell lines, CIMIC similarly identified Fun-CIM and Dys-CIM trajectories, with directionally conserved induction of immunostimulatory and inflammatory programs, including viral mimicry, antigen presentation, and inflammatory cell death, in Fun-CIM cell lines and proteostatic and mitochondrial stress-adaptation programs in Dys-CIM cell lines. These findings indicate that tumor-cell responses contribute to, but do not fully define, the broader CIM trajectory observed in patient tumors, thus suggesting that CIM is not solely a product of chemotherapy-induced remodeling of the immune microenvironment, but also reflects tumor-intrinsic responses that may shape the subsequent immunologic consequences of therapy. This provides a potential framework for understanding why otherwise similar chemotherapeutic exposures can produce divergent immune outcomes and raises the possibility that tumor-intrinsic stress-adaptation programs could be therapeutically targeted to redirect CIM toward productive antitumoral immunity (20,30).

Although both trajectories were defined among tumors with residual disease, they exhibited divergent chemotherapy-induced remodeling that was not consistently distinguishable from baseline transcriptional state. Dys-CIM tumors exhibited greater baseline immune enrichment, immune-related transcriptional programs, and checkpoint expression but did not preferentially induce adaptive immune programs and subsequently demonstrated contraction of cytotoxic immune compartments. Conversely, Fun-CIM tumors were less immune enriched at baseline but preferentially induced adaptive immune activation, antigen presentation, and inflammatory signaling. These findings suggest that baseline TME activity and composition, including TIL abundance, a widely used predictive biomarker, may not always reliably predict whether a tumor will mount a productive immunostimulatory response to chemotherapy. To directly assess whether CIM induction was intrinsically encoded by baseline transcriptional state, we evaluated the correlation between pre-treatment CIM gene-expression profiles and corresponding ΔGE profiles. These correlations were weak across the patient cohorts and TNBC cell lines (**Figure SM1**). Thus, CIM trajectories reflect chemotherapy-induced transcriptional remodeling that is not simply determined by baseline expression. Instead, how tumors transcriptionally respond to chemotherapy-induced stress may be more critical in determining the resulting CIM trajectory. This notion aligns with observations from the SMC cohort, in which baseline immune features were associated with early on-treatment response, but on-treatment features and TILs localization were ultimately more informative than baseline features alone for predicting clinical outcome (21).

CIM trajectory-specific genes also showed opposing clinical associations at both the gene and signature levels. In NKI, significant associations between treatment-induced ΔGE were predominantly favorable for Fun-CIM and adverse for Dys-CIM; consistently, increased Fun-CIM ΔssGSEA was associated with improved RFS, whereas increased Dys-CIM ΔssGSEA was significantly associated with worse RFS. However, the small cohort (N = 20; six events) and lack of external longitudinal validation limit these findings to exploratory prognostic associations. Baseline analysis in the chemotherapy-treated METABRIC (N = 412) and SCAN-B (N = 2,462) cohorts further showed favorable associations for the Fun-CIM signature and adverse associations for the Dys-CIM signature after adjustment for clinicopathological variables. Although baseline analysis cannot validate the prognostic effect of chemotherapy-induced changes, the reproducible opposing associations across the independent cohorts further support the clinical relevance of genes associated with and potentially defining CIM-trajectories. Nonetheless, the full prognostic impact of CIM may be best captured through longitudinal assessment of treatment-induced changes.

From a translational perspective, CIMIC provides a foundation for two clinically motivated research directions: first, identifying baseline molecular or genetic features that predict which CIM trajectory a patient will follow in response to a particular agent (baseline CIM trajectory prediction), and second, identifying potential pharmacologic strategies that redirect Dys-CIM tumors toward Fun-CIM programming (therapeutic reprogramming). Future studies applying CIMIC to larger paired pre- and post-treatment datasets may uncover additional, potentially latent CIM trajectories and refine CIM trajectory characterizations while generating the data needed to develop models that predict CIM trajectory from baseline tumor features. In parallel, mechanistic studies of the proteostatic and mitochondrial stress programs enriched in Dys-CIM may identify pharmacologic vulnerabilities whose inhibition could disrupt stress-adaptive responses and promote productive CIM (59,60).

Several limitations warrant consideration. Transcriptomic data from pre- and on-treatment biopsies of pCR-achieving tumors are limited, restricting our analyses primarily to pre- and post-treatment specimens and to patients with residual disease. In addition, our data sets are limited by a small sample size, heterogeneous NACT regimens, and variable treatment duration. While these limitations are inherent to longitudinal neoadjuvant datasets, it nonetheless limits further investigation of subtype-, regimen-, and time-dependent determinants of CIM induction. Importantly, our orthogonal *in vitro* analyses partially address these constraints by demonstrating consistent Fun-CIM and Dys-CIM trajectory assignment under controlled chemotherapeutic perturbation, supporting the reproducibility of CIMIC-derived classifications across experimental contexts. However, these cell line models isolate tumor-intrinsic responses and do not capture tumor-immune or stromal interactions that contribute to CIM *in vivo*. Moreover, the use of bulk transcriptomic data precludes definitive attribution of CIM programs to specific cellular compartments in patient tumors, and immune deconvolution analyses provide only inferential estimates of TME composition. Future application of CIMIC to more physiologically relevant three-dimensional patient-derived tumor-immune coculture models may help resolve the contribution and interaction of tumor, immune, and stromal compartments to divergent CIM trajectories. Additionally, prognostic analyses in the independent METABRIC and SCAN-B cohorts relied on baseline expression of CIM trajectory-associated signatures rather than treatment-induced ΔGE and therefore do not directly assess the clinical relevance of the CIM trajectories according to their dynamic nature. Finally, the binary Fun-CIM/Dys-CIM framework, while robust in current datasets, likely represents a simplification of a more continuous and context-dependent landscape of chemoimmunomodulatory responses. Future studies applying CIMIC to larger cohorts may enable refinement the current binary Fun-CIM and Dys-CIM framework into a more granular continuum of chemotherapy-induced immunomodulatory responses. Nonetheless, our findings collectively establish CIM as a dynamic and clinically consequential process in breast cancer. Furthermore, CIMIC provides a framework for systematically understanding and classifying CIM trajectories, which can be leveraged in future studies to identify baseline predictors of CIM, elucidate mechanisms driving divergent trajectories, and inform strategies to maximize beneficial immunomodulation.

## Author Contributions

I.L., K.H.S., E.M., M.M., K.L.C., N.S., and M.O.G wrote the manuscript; I.L. and M.O.G contributed to the conceptualization; I.L. and M.O.G designed the research; I.L. and M.O.G developed the software; E.M., K.H.S., K.L.C., M.M., and M.O.G performed the research; I.L., E.M., and M.O.G. analyzed the data. M.O.G funded the work.

I.L.: Conceptualization; Methodology; Software; Investigation; Formal analysis; Writing, original draft; Writing, review and editing.

E.M.: Conceptualization; Designing and performing research; Investigation; Formal analysis; Writing, original draft; Writing, review and editing.

K.H.S.: Designing and performing research; Writing, review and editing;

M.M.: Designing and performing research; Writing, original draft; Writing, review and editing.

K.L.C.: Designing and performing research; Conceptualization; Writing, original draft; Writing, review and editing.

N.S.: Writing, original draft; Writing, review and editing.

M.O.G.: Conceptualization; Methodology; Supervision; Funding acquisition; Project administration; Writing, original draft; Writing, review and editing.

## Disclosure Statement

The authors have no relevant affiliations or financial involvement with any organization or entity with a financial interest in or financial conflict with the subject matter or materials discussed in the manuscript. This includes employment, consultancies, honoraria, stock ownership or options, expert testimony, grants or patents received or pending, or royalties.

## Data Sharing Statement

The CIMIC pipeline, alongside code for figure generation are readily available for use at https://github.com/Gbadamosi-Lab/CIMIC. Gene Expression Data used for this project is publicly available via GEO Omnibus under the following accession numbers: GSE191127 (NKI), GSE123845 (SMC), GSE122630 (NEO), and GSE345047 (TNBC Cell Lines).

## SUPPLEMENTAL FIGURES AND FIGURE LEGENDS

**Supplemental Figure S1.**
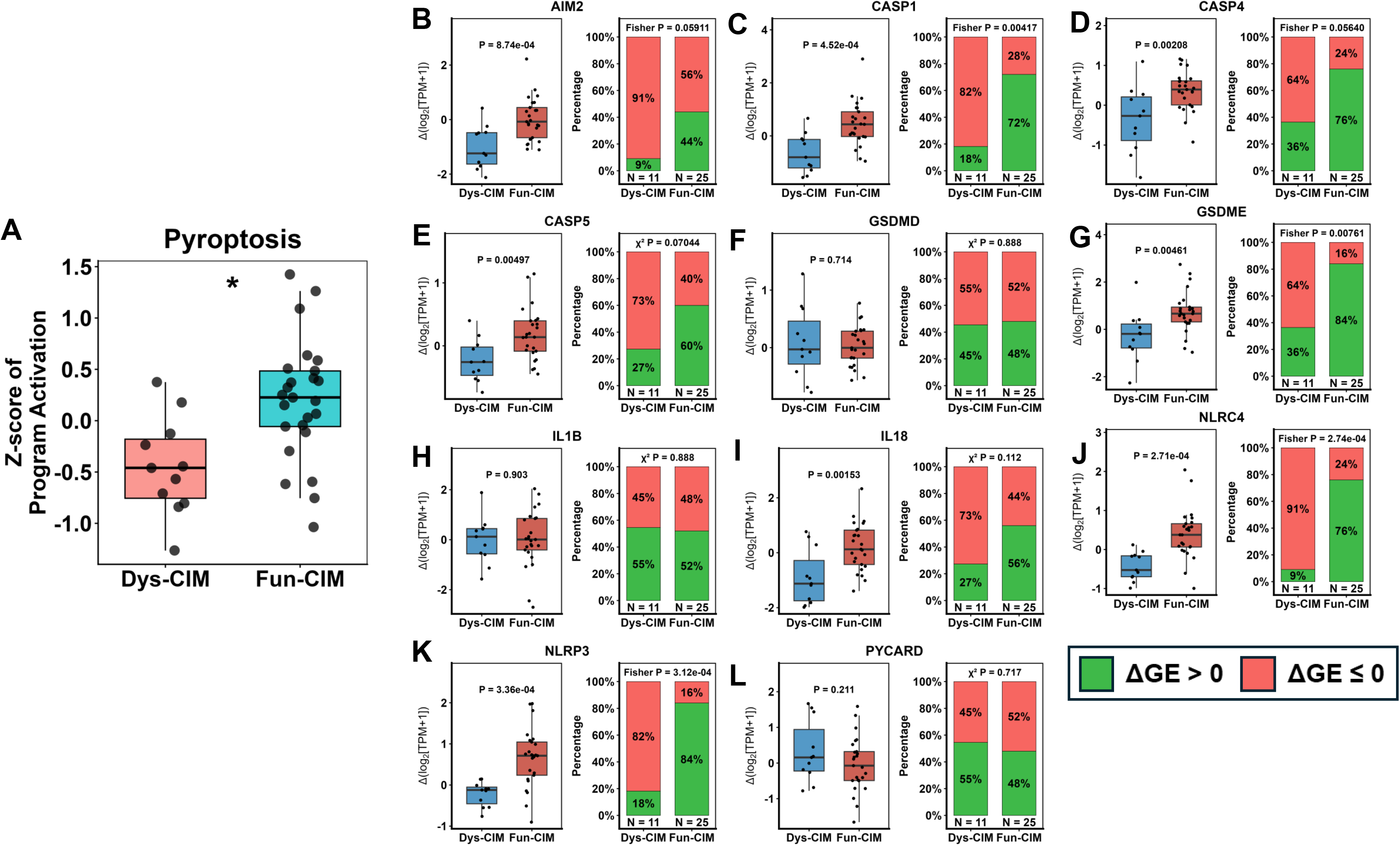
Differential induction of pyroptotic cell death program. **(A)** Boxplot depicts the relative differential induction (Z-score of average ΔGE values) of pyroptotic cell death program as in **Figure 2F** in Fun-CIM (blue) and Dys-CIM (red) tumors. Statistically significant differences for FDR-adjusted p values are indicated as follows: * ≤ 0.05; ** ≤ 0.01; *** ≤ 0.005; **** ≤ 0.001. **(B-L)** Boxplots display differential gene induction (Δlog2[TPM+1]) in Dys-CIM and Fun-CIM tumors for genes utilized in single-sample gene set analysis, including *AIM2*, *CASP1*, *CASP4*, *CASP5*, *GSDMD*, *GSDME*, *IL1B*, *IL18*, *NLRC4*, *NLRP3*, and *PYCARD.* Stacked bar plots show the proportion of tumors exhibiting induction (green) versus no induction (red) within each trajectory group. p *value*s were computed using the Wilcoxon rank-sum test and Fisher’s exact or chi-squared test, as indicated, for comparison of average induction values and proportion of tumors exhibiting induction, respectively.

**Supplemental Figure S2.**
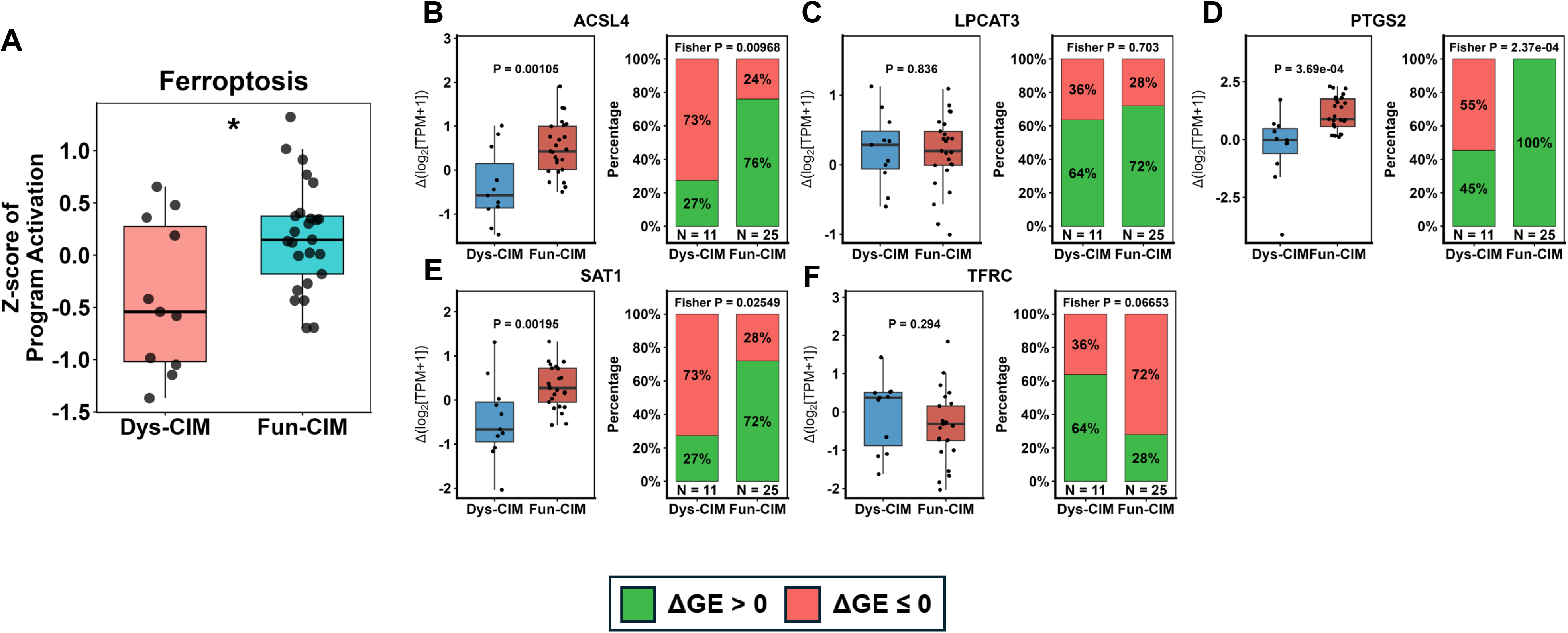
Differential induction of ferroptotic cell death program. **(A)** Boxplot depicts the relative differential induction (Z-score of average ΔGE values) of ferroptotic cell death program as in **Figure 2G** in Fun-CIM (blue) and Dys-CIM (red) tumors. Statistically significant differences for FDR-adjusted p values are indicated as follows: * ≤ 0.05; ** ≤ 0.01; *** ≤ 0.005; **** ≤ 0.001. **(B-F)** Boxplots display differential gene induction (Δlog2[TPM+1]) in Dys-CIM and Fun-CIM tumors for genes utilized in single-sample gene set analysis, including *ACSL4, LPCAT3, PTGS2, SAT1, and TFRC.* Stacked bar plots show the proportion of tumors exhibiting induction (green) versus no induction (red) within each trajectory group. p values were computed using the Wilcoxon rank-sum test and Fisher’s exact or chi-squared test for comparison of average induction values and proportion of tumors exhibiting induction, respectively.

**Supplemental Figure S3.**
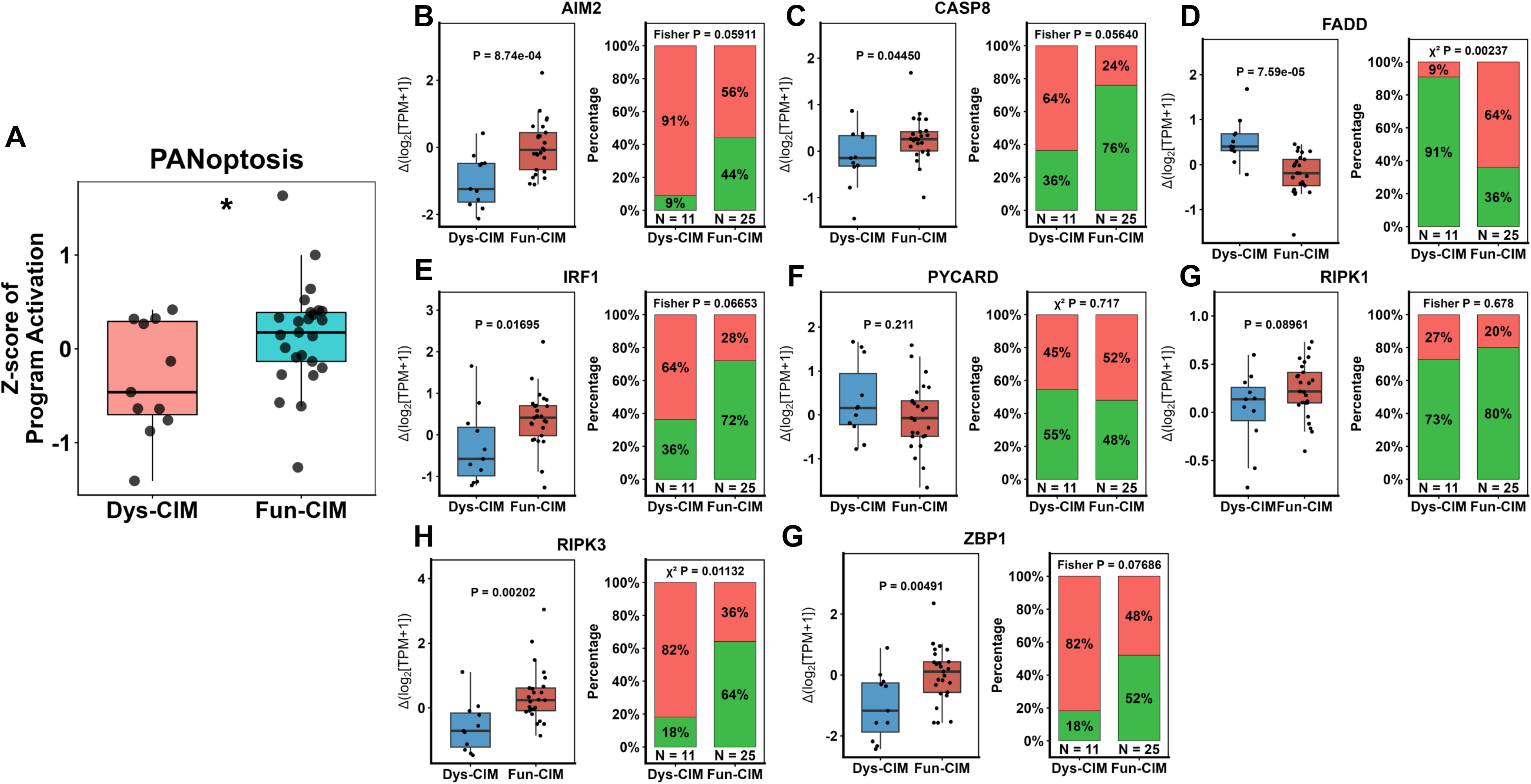
Differential induction of PANoptotic cell death program. **(A)** Boxplot depicts the relative differential induction (Z-score of average ΔGE values) of PANoptotic cell death program as in **Figure 2H** in Fun-CIM (blue) and Dys-CIM (red) tumors. Statistically significant differences for FDR-adjusted p values are indicated as follows: * ≤ 0.05; ** ≤ 0.01; *** ≤ 0.005; **** ≤ 0.001. **(B-I)** Boxplots display differential gene induction (Δlog2[TPM+1]) in Dys-CIM and Fun-CIM tumors for genes utilized in single-sample gene set analysis, including *AIM2, CASP8, FADD, IRF1, RIPK1, RIPK3, PYCARD, and ZBP1.* Stacked bar plots show the proportion of tumors exhibiting induction (green) versus no induction (red) within each trajectory group. p values were computed using the Wilcoxon rank-sum test and Fisher’s exact or chi-squared test for comparison of average induction values and proportion of tumors exhibiting induction, respectively.

**Supplemental Figure S4.**
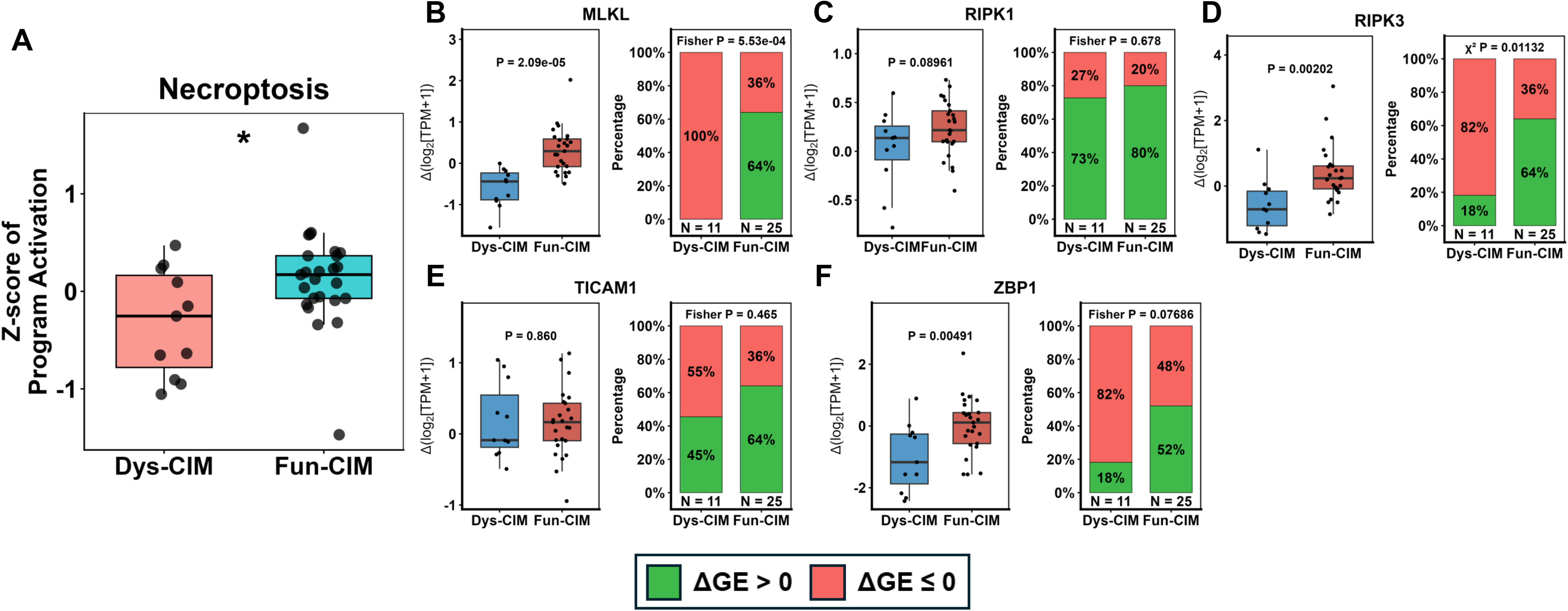
Differential induction of necroptotic cell death program. **(A)** Boxplot depicts the relative differential induction (Z-score of average ΔGE values) of necroptotic cell death program as in **Figure 2I** in Fun-CIM (blue) and Dys-CIM (red) tumors. Statistically significant differences for FDR-adjusted p values are indicated as follows: * ≤ 0.05; ** ≤ 0.01; *** ≤ 0.005; **** ≤ 0.001. **(B-F)** Boxplots display differential gene induction (Δlog2[TPM+1]) in Dys-CIM and Fun-CIM tumors for genes utilized in single-sample geneset analysis, including *MLKL*, *RIPK1*, *RIPK3*, *TICAM1*, and *ZBP1.* Stacked bar plots show the proportion of tumors exhibiting induction (green) versus no induction (red) within each trajectory group. p values were computed using the Wilcoxon rank-sum test and Fisher’s exact or chi-squared test for comparison of average induction values and proportion of tumors exhibiting induction, respectively.

**Supplemental Figure S5.**
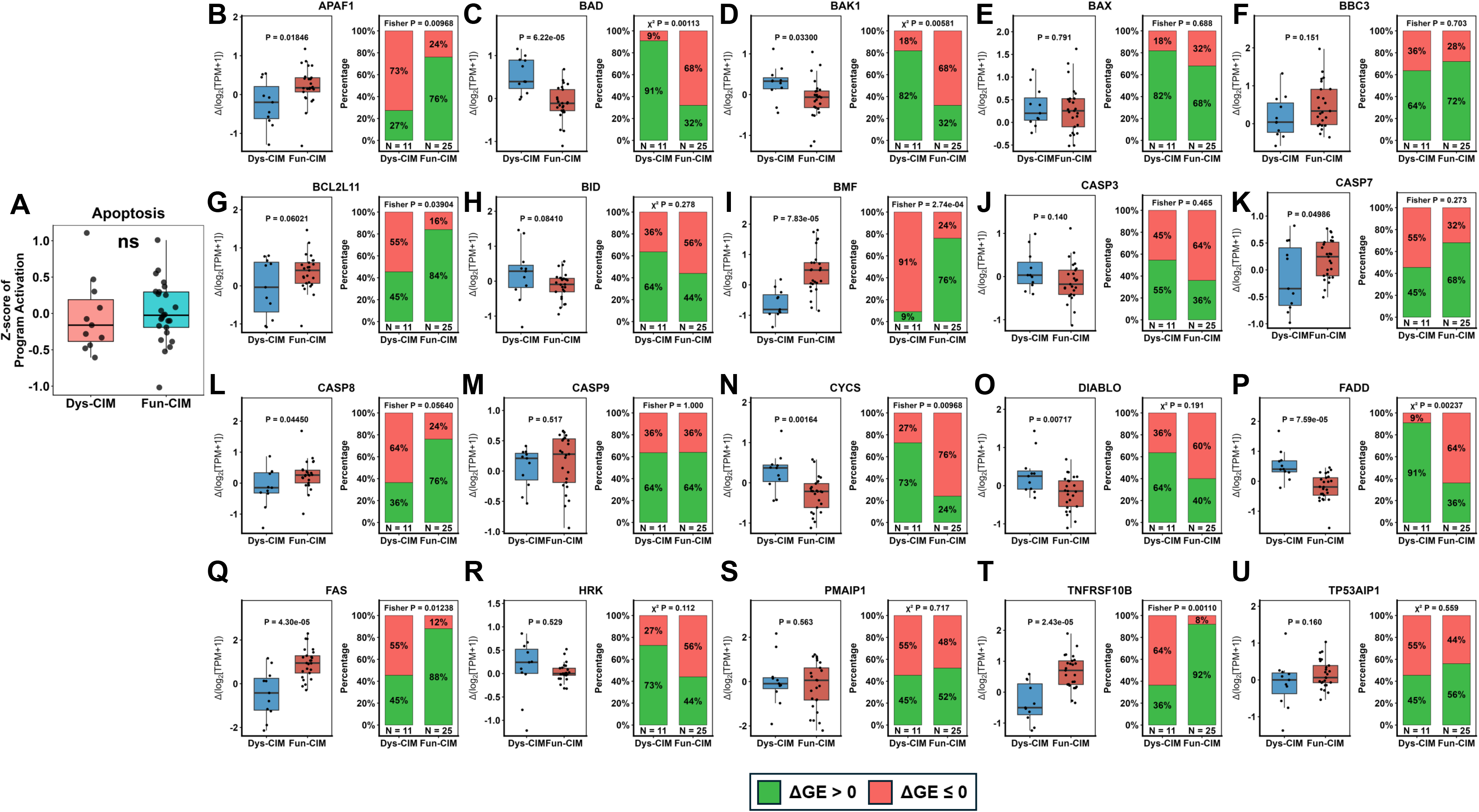
Differential induction of apoptotic cell death program. **(A)** Boxplot depicts the relative differential induction (Z-score of average ΔGE values) of apoptotic cell death program as in **Figure 2J** in Fun-CIM (blue) and Dys-CIM (red) tumors. Statistically significant differences for FDR-adjusted p values are indicated as follows: * ≤ 0.05; ** ≤ 0.01; *** ≤ 0.005; **** ≤ 0.001. **(B-U)** Boxplots display differential gene induction (Δlog2[TPM+1]) in Dys-CIM and Fun-CIM tumors for genes utilized in single-sample geneset analysis, including *APAF1, BAD, BAK1, BAX, BBC3, BCL2L11, BID, BMF, CASP3, CASP7, CASP8, CASP9, CYCS, DIABLO, FADD, FAS, HRK, PMAIP1, TNFRSF10B, and TP53AIP1.* Stacked bar plots show the proportion of tumors exhibiting induction (green) versus no induction (red) within each trajectory group. p values were computed using the Wilcoxon rank-sum test and using Fisher’s exact or chi-squared test, as indicated, for comparison of average induction values and proportion of tumors exhibiting induction, respectively.

**Supplemental Figure S6.**
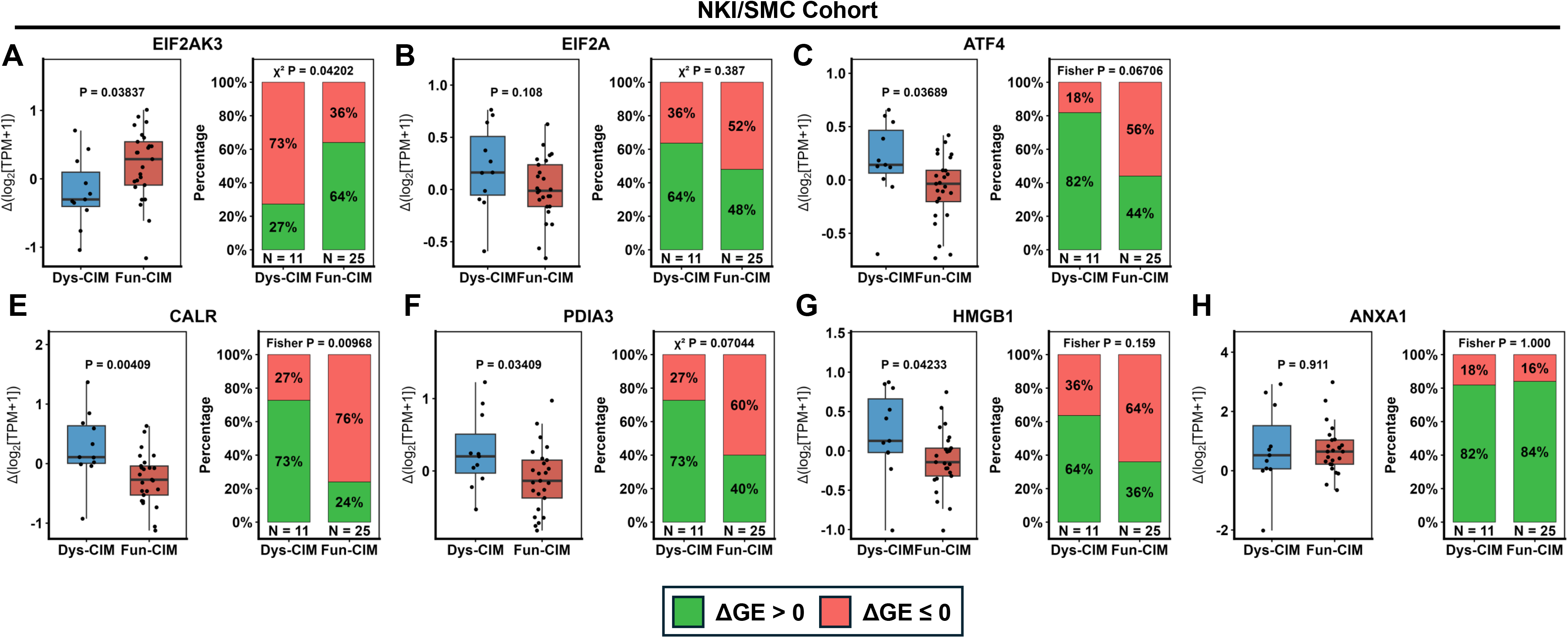
Differential induction of ICD genes across CIM trajectories. (A-H) Boxplots display differential gene induction (Δlog2[TPM+1]) in Dys-CIM and Fun-CIM tumors *EIF2AK3, EIF2A, ATF4, CALR, PDIA3, HMGB1, and ANXA1*. Stacked bar plots show the proportion of tumors exhibiting induction (green) versus no induction (red) within each trajectory group. Boxed legend applies to adjacent stacked bar plots indicating percentage of samples with positive induction (ΔGE > 0) or negative/no induction (ΔGE ≤ 0). p values were computed using the Wilcoxon rank-sum test and using Fisher’s exact or chi-squared test, as indicated, for comparison of average induction values and proportion of tumors exhibiting induction, respectively.

**Supplemental Figure S7.**
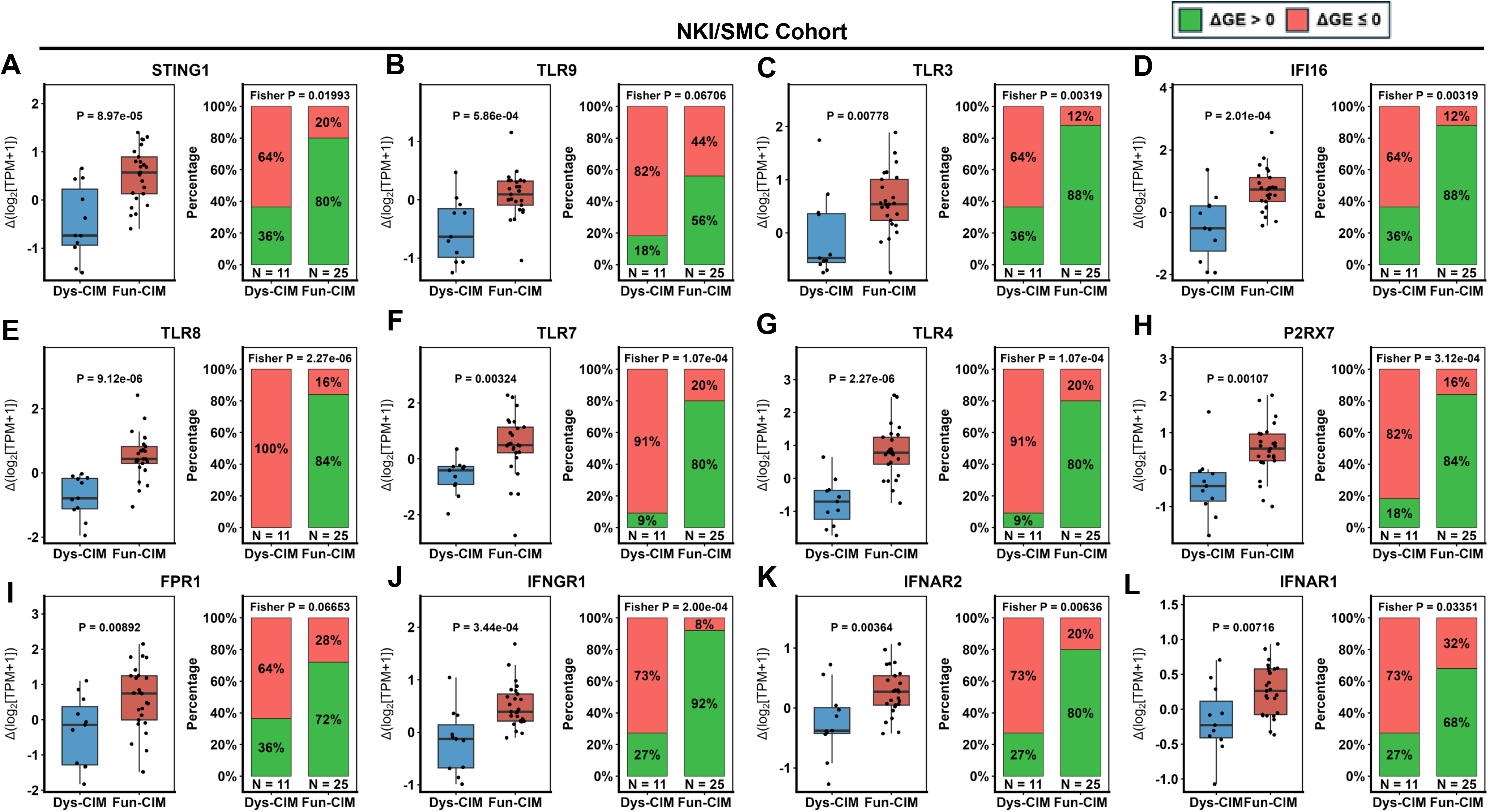
Fun-CIM tumors have greater induction of CIM sensors. (A-H) Boxplots display differential gene induction (Δlog2[TPM+1]) in Dys-CIM (blue) and Fun-CIM (red) tumors for key innate immune sensing and interferon signaling genes, including *IFI16*, *TLR9*, *TLR3*, *TLR7*, *TLR8*, *P2RX7*, *FPR1*, *TLR4*, *IFNAR1*, *IFNAR2*, *IFNGR1*, and *STING1*. Stacked bar plots show the proportion of tumors exhibiting induction (green) versus no induction (red) within each trajectory group. Boxed legend applies to stacked barplots indicating the percentage of samples with positive induction (ΔGE > 0) or negative/no induction (ΔGE ≤ 0). p values were computed using the Wilcoxon rank-sum test and using Fisher’s exact test for comparison of average induction values and proportion of tumors exhibiting induction, respectively.

**Supplemental Figure S8.**
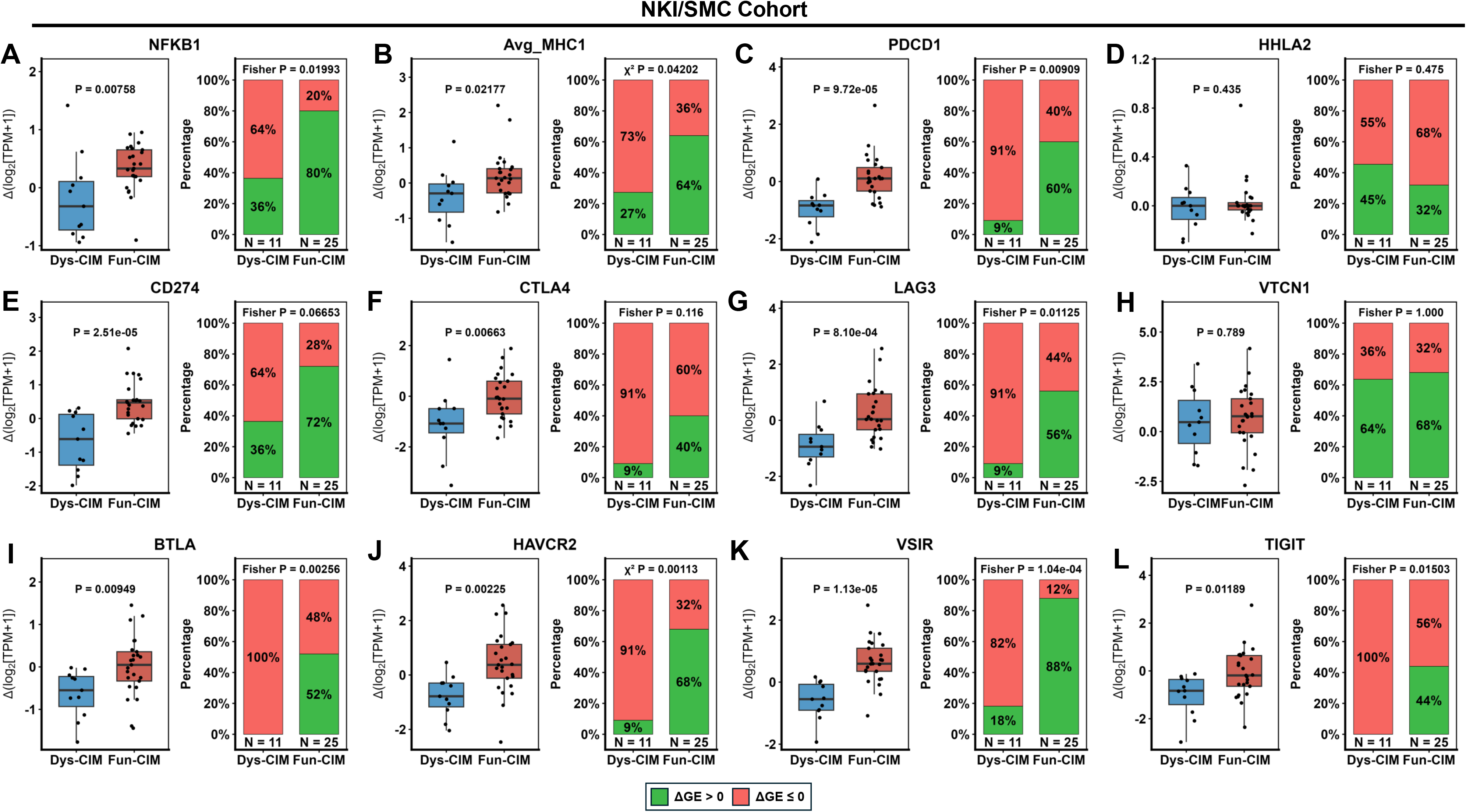
Differential induction of antigen-presentation genes and immune checkpoint-related genes across CIM trajectories. (A–L) Boxplots show differential gene induction (Δlog2[TPM+1]) between pre- and post-chemotherapy samples in Dys-CIM and Fun-CIM tumors for *NFKB1* (A), Avg_MHC1 (average expression of *HLA-A, HLA-B,* and *HLA-C*) **(B)**, *PDCD1* (C), *HHLA2* (D), *CD274* (PD-L1) **(E)**, *CTLA4* (F), *LAG3* (G), *VTCN1* (H), *BTLA* (I), *HAVCR2* (J), *VSIR* (K), and *TIGIT* (L). Adjacent stacked bar plots show the proportion of tumors exhibiting positive induction (ΔGE > 0; green) versus no or negative induction (ΔGE ≤ 0; red) within each CIM trajectory. Fun-CIM tumors exhibited greater induction of multiple antigen-presentation and immune checkpoint-related genes compared with Dys-CIM tumors. p values above boxplots indicate comparisons of ΔGE between CIM trajectories using the Wilcoxon rank-sum test, whereas p values above stacked bar plots indicate comparisons of the proportions of tumors exhibiting positive versus no/negative induction using Fisher’s exact or chi-squared test, as indicated.

**Supplemental Figure S9.**
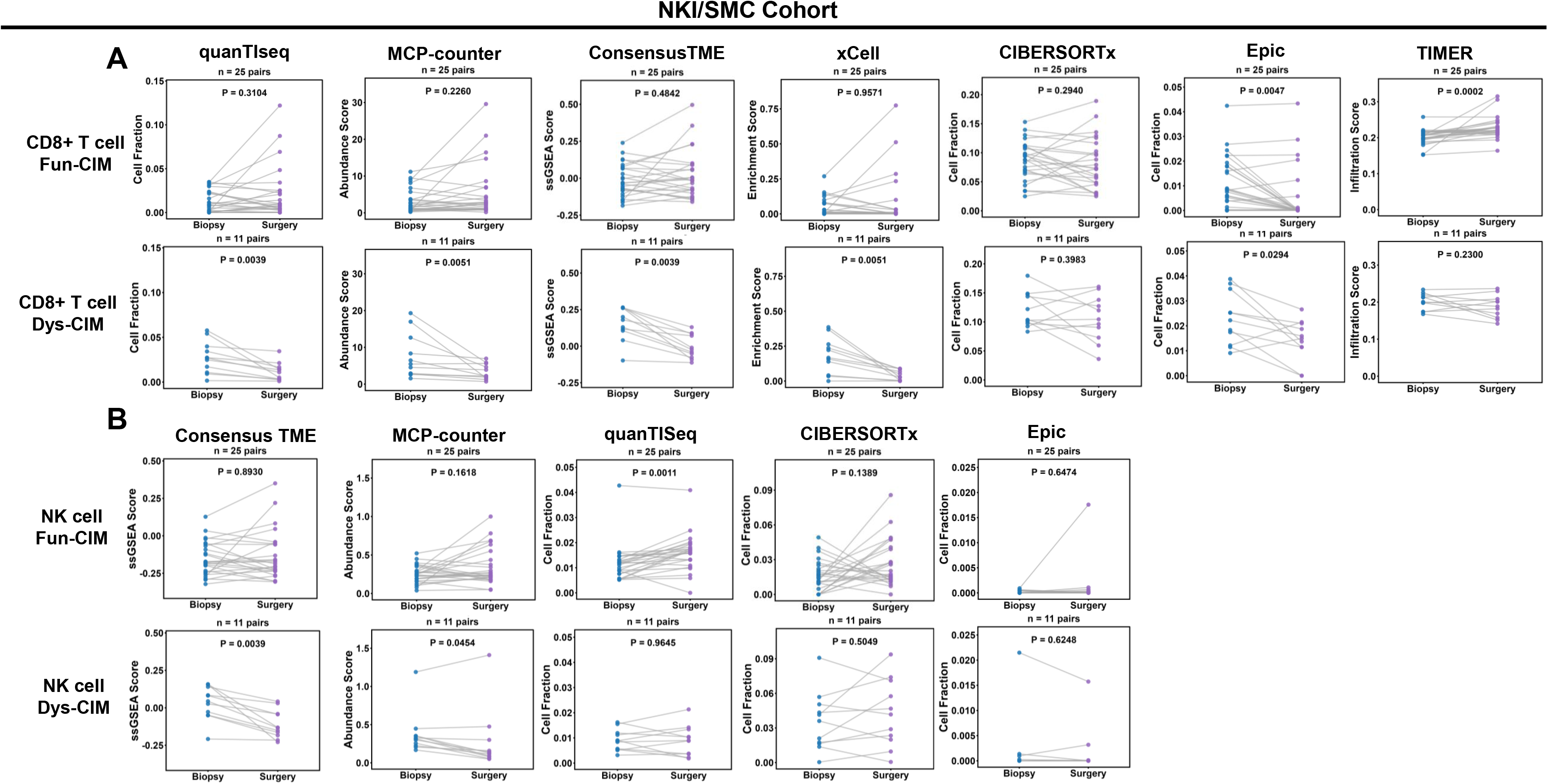
Biopsy-to-surgery changes in CD8⁺ T-cell abundance across CIM trajectories in NKI/SMC cohorts. Paired comparisons of immune-cell abundance between pre-treatment biopsy and post-treatment surgical specimens in Fun-CIM tumors (n = 25) and Dys-CIM tumors (n = 11). **(A)** CD8⁺ T-cell abundance estimated using quanTIseq, MCP-counter, ConsensusTME, xCell, CIBERSORTx, EPIC, and TIMER. **(B)** NK-cell abundance estimated using ConsensusTME, MCP-counter, quanTIseq, CIBERSORTx, and EPIC. Each dot represents an individual tumor sample, with lines connecting matched biopsy and surgical specimens. Dys-CIM tumors exhibited significant reductions in CD8⁺ T-cell abundance following chemotherapy across quanTIseq, MCP-counter, ConsensusTME, xCell, and EPIC methods. NK-cell abundance was also significantly reduced in Dys-CIM tumors by ConsensusTME and MCP-counter, whereas no significant overall reduction was observed in Fun-CIM tumors, with a subset of patients showing increased in CD8⁺ T-cell and NK-cell abundance following treatment. Displayed p *value*s were calculated using two-sided paired Wilcoxon signed-rank tests.

**Supplemental Figure S10.**
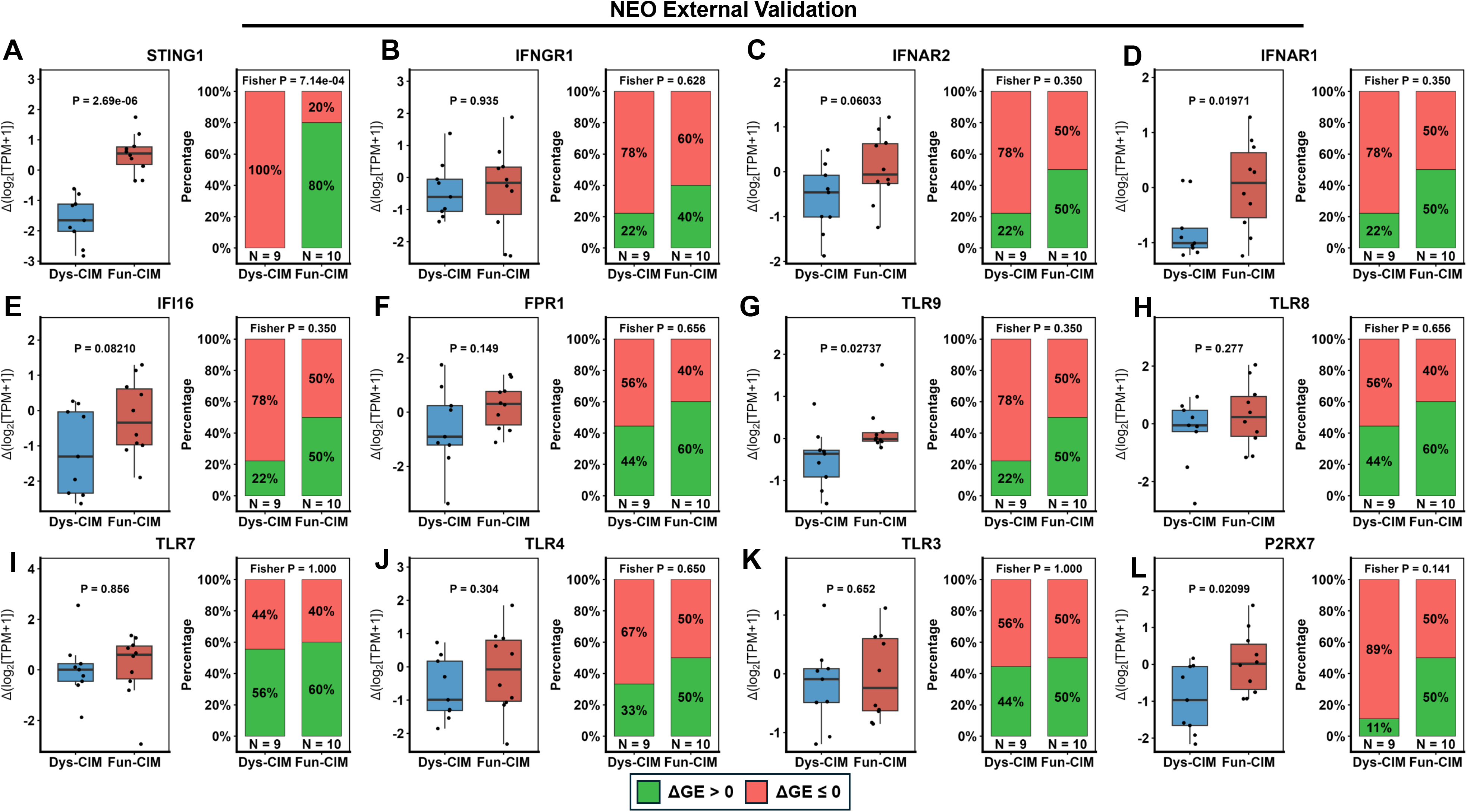
Validation of enhanced CIM sensor induction in Fun-CIM tumors in the NEO cohort. (A–L) Boxplots show differential gene induction (Δlog2[TPM+1]) between pre- and post-chemotherapy samples in Dys-CIM (blue) and Fun-CIM (red) tumors for key innate immune sensing and interferon signaling genes, including *STING1* (A), *IFNGR1* (B), *IFNAR2* (C), *IFNAR1* (D), *IFI16* (E), *FPR1* (F), *TLR9* (G), *TLR8* (H), *TLR7* (I), *TLR4* (J), *TLR3* (K), and *P2RX7* (L). Adjacent stacked bar plots show the proportion of tumors exhibiting induction (green) versus no induction (red) within each trajectory group. Boxed legend applies to stacked barplots indicating percentage of samples with positive induction (ΔGE > 0) or negative/no induction (ΔGE ≤ 0) p values were computed using the Wilcoxon rank-sum test and Fisher’s exact test for comparison of average induction values and proportion of tumors exhibiting induction, respectively.

**Supplemental Figure S11.**
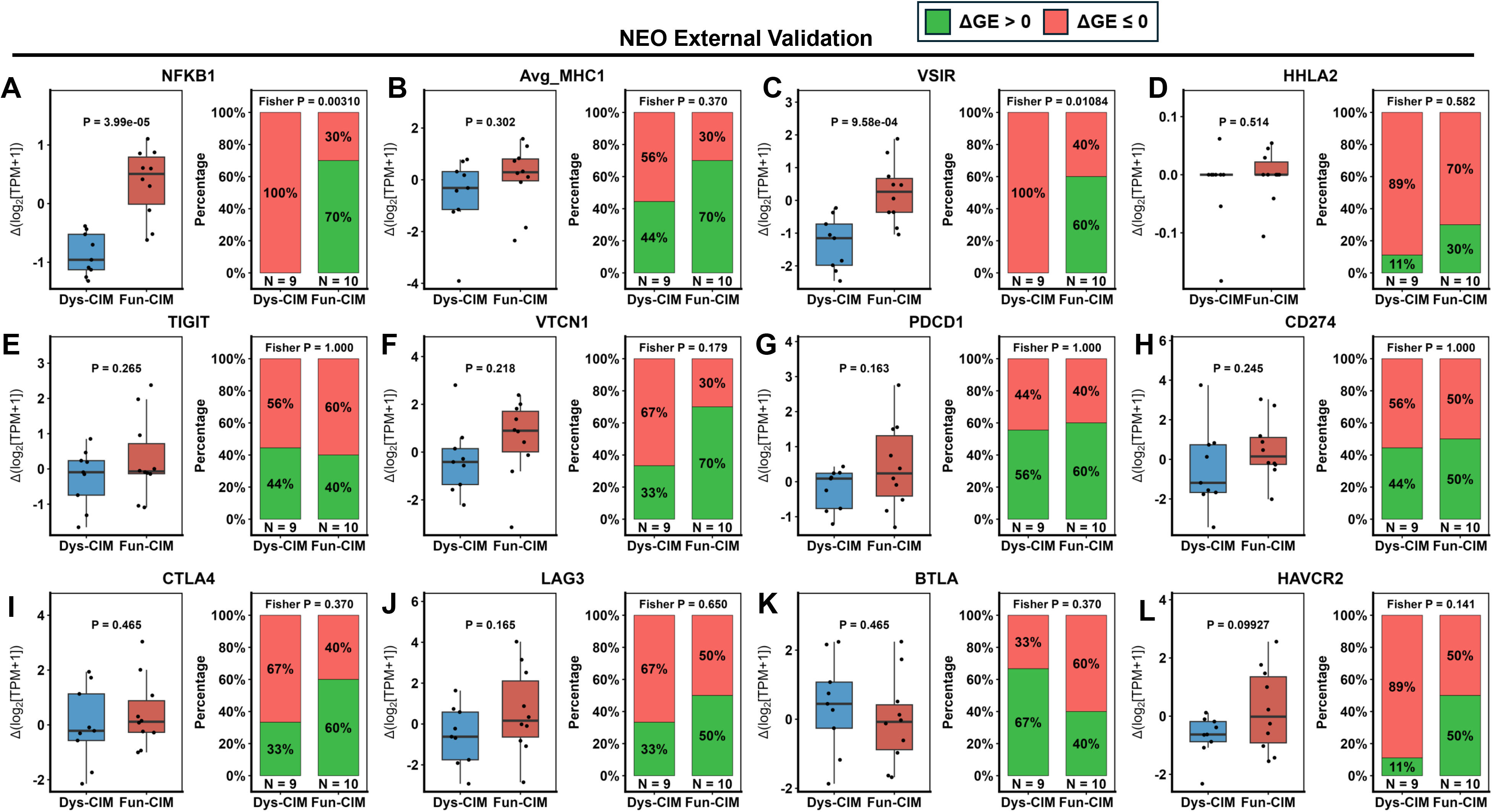
Induction of inflammatory signaling, antigen-presentation, and immune checkpoint-related genes across CIM trajectories in the NEO cohort. (A–L) Boxplots show differential gene induction (Δlog2[TPM+1]) between pre- and post-chemotherapy samples in Dys-CIM (blue) and Fun-CIM (red) tumors for *NFKB1* (A), Avg_MHC1 (average expression of *HLA-A, HLA-B,* and *HLA-C*) (B), *VSIR* (C), *HHLA2* (D), *TIGIT* (E), *VTCN1* (F), *PDCD1* (G), *CD274* (PD-L1) (H), *CTLA4* (I), *LAG3* (J), *BTLA* (K), and *HAVCR2* (L). Adjacent stacked bar plots show the proportion of tumors exhibiting positive induction (ΔGE > 0; green) versus no or negative induction (ΔGE ≤ 0; red) within each CIM trajectory. Fun-CIM tumors exhibited significantly greater induction of *NFKB1* and *VSIR*, with a significantly higher proportion of Fun-CIM tumors exhibiting positive induction of both genes. p values above boxplots indicate comparisons of ΔGE between CIM trajectories using the Wilcoxon rank-sum test, whereas p values above stacked bar plots indicate comparisons of the proportions of tumors exhibiting positive versus no/negative induction using Fisher’s exact test.

**Supplemental Figure S12.**
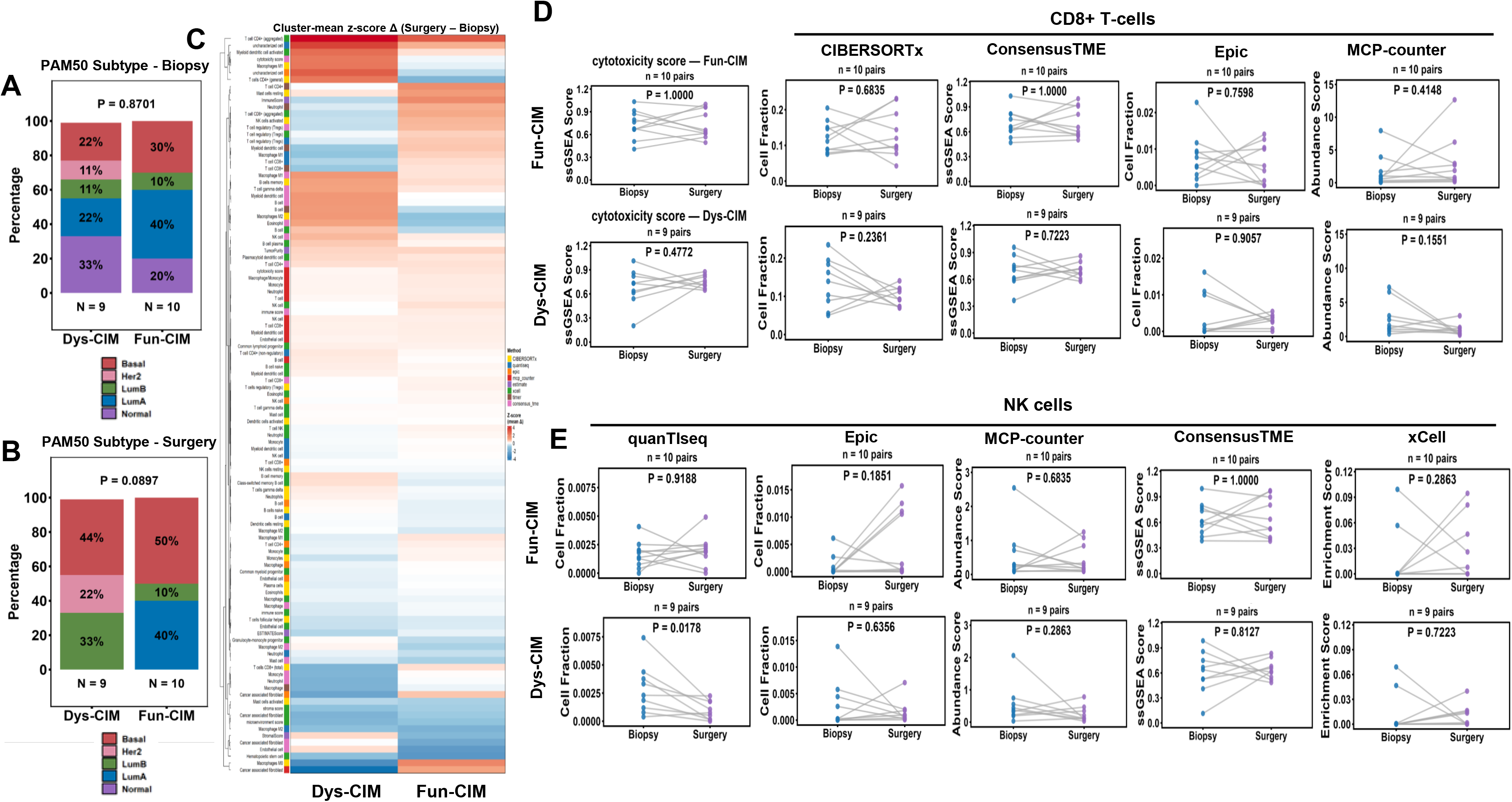
PAM50 subtyping and deconvolution-based immune profiling by CIMIC derived trajectories in the NEO cohort. Stacked bar plots showing the distributions of PAM50 molecular subtypes in Dys-CIM and Fun-CIM tumors at pretreatment biopsy **(A)** and post-treatment surgery **(B)**. Displayed *p*-values were calculated using Fisher’s exact test. **(C)** Heatmap summarizing paired biopsy-to-surgery changes in immune-cell abundance and immune-related scores estimated using eight deconvolution methods. Changes were calculated as surgery minus biopsy and standardized as Z scores. Rows represent immune-cell types or immune scores, and columns represent the mean Z-scored change within each CIM trajectory; red and blue indicate relative increases and decreases, respectively. **(D)** Paired biopsy-to-surgery changes in ConsensusTME cytotoxicity scores and CD8⁺ T-cell abundance estimated using CIBERSORTx, ConsensusTME, EPIC, and MCP-counter. **(E)** Paired changes in NK-cell abundance estimated using quanTIseq, EPIC, MCP-counter, ConsensusTME, and xCell. Each dot represents an individual tumor sample, with lines connecting matched biopsy and surgical specimens. Displayed *p*-values for paired comparisons were calculated using two-sided Wilcoxon signed-rank tests.

**Supplemental Figure S13.**
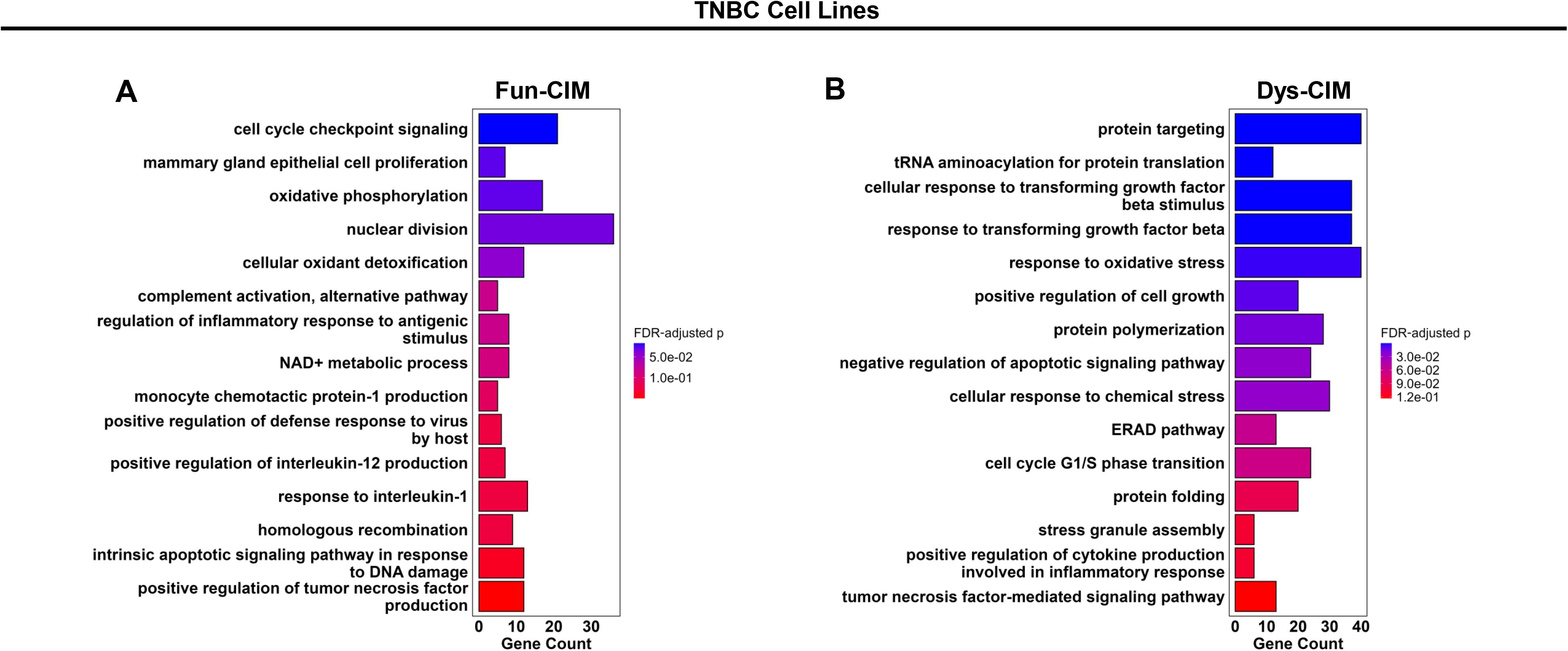
Fun-CIM and Dys-CIM TNBC cell lines exhibit divergent transcriptional programs induced by chemotherapy. Top 15 enriched Gene Ontology (GO) biological processes among genes preferentially induced in Fun-CIM (A) and Dys-CIM (B) cell lines following 48 hours of epirubicin treatment. Bar length represents the number of genes overlapping each GO biological process, and color indicates the FDR-adjusted p value.

**Supplemental Figure S14.**
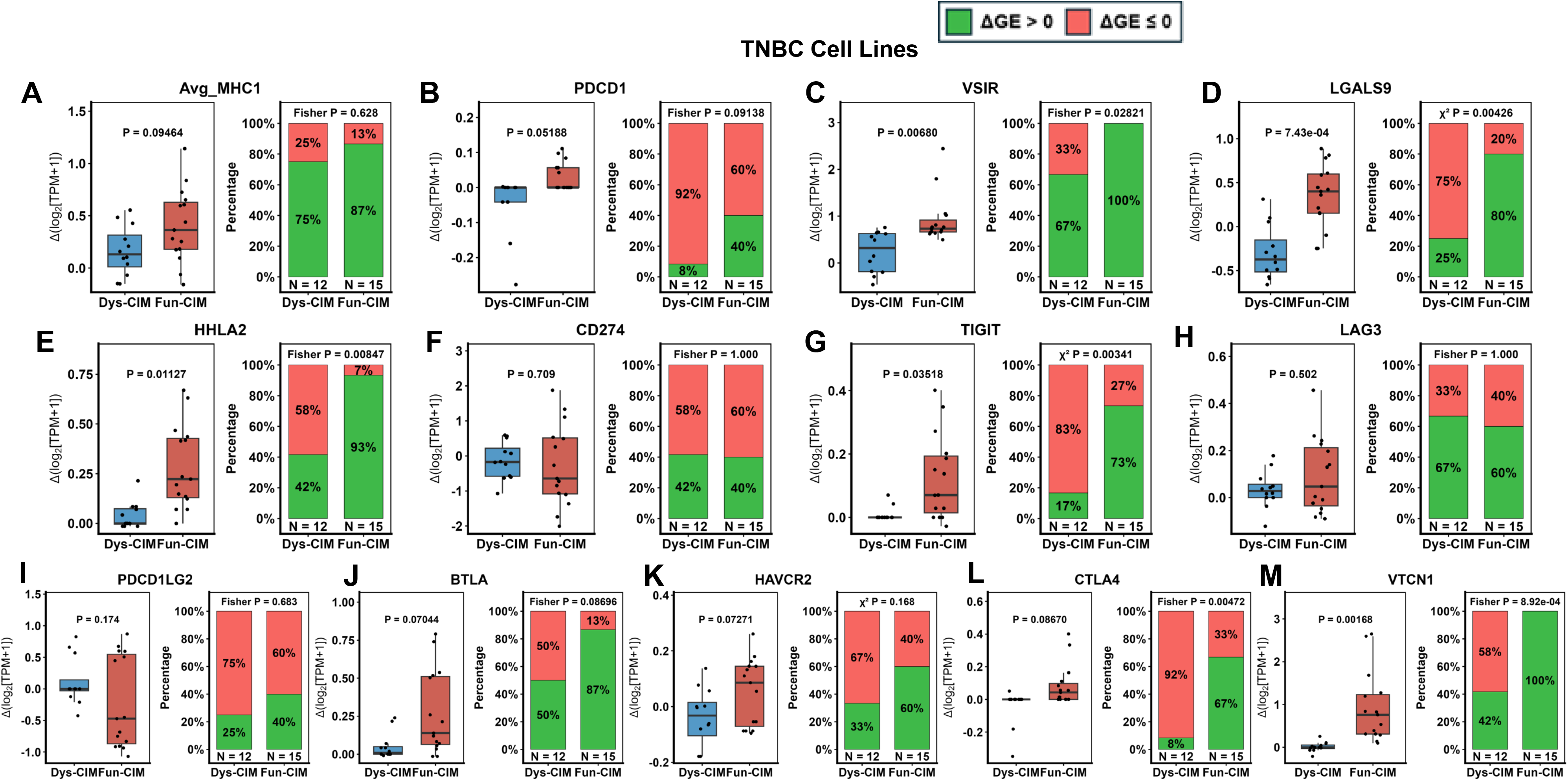
Induction of immune checkpoints in TNBC cell lines. (A-H) (A–M) Boxplots show differential gene induction (Δlog2[TPM+1]) following epirubicin treatment in Dys-CIM (blue) and Fun-CIM (red) cell lines for Avg_MHC1 (average expression of *HLA-A, HLA-B,* and *HLA-C*) **(A)**, *PDCD1* **(B)**, *VSIR* **(C)**, *LGALS9* **(D)**, *HHLA2* **(E)**, *CD274* (PD-L1) **(F)**, *TIGIT* **(G)**, *LAG3* **(H)**, *PDCD1LG2* (PD-L2) **(I)**, *BTLA* **(J)**, *HAVCR2* **(K)**, *CTLA4* **(L)**, and *VTCN1* **(M)**. Adjacent stacked bar plots show the proportion of samples exhibiting positive induction (ΔGE > 0; green) versus no or negative induction (ΔGE ≤ 0; red) within each CIM trajectory. Fun-CIM cell lines exhibited greater induction of several immune checkpoint-related genes, including *VSIR, LGALS9, HHLA2, TIGIT,* and *VTCN1*. p values above boxplots indicate comparisons of ΔGE between CIM trajectories using the Wilcoxon rank-sum test, whereas p values above stacked bar plots indicate comparisons of the proportions of samples exhibiting positive versus no/negative induction using Fisher’s exact or chi-squared test, as indicated.

**Supplemental Figure S15.**
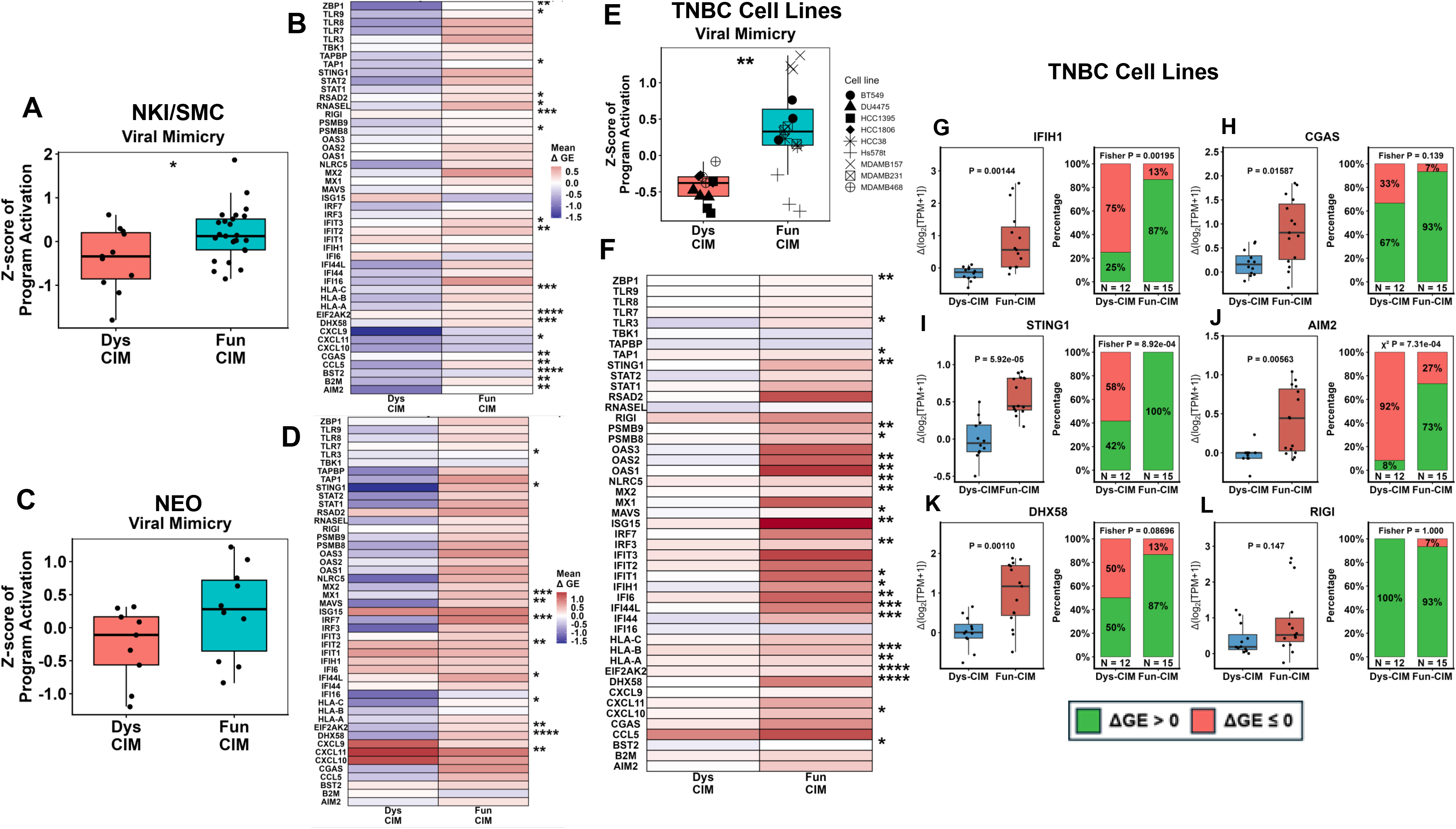
Fun-CIM trajectories exhibit greater induction of viral mimicry programs and cytosolic nucleic acid sensors in response to chemotherapy across tumors and cell line datasets. (A–F) Relative differential induction of a 47-gene viral mimicry signature in Fun-CIM and Dys-CIM tumors from the NKI/SMC and NEO cohorts and in TNBC cell lines. (A) Box plot depicts the relative differential induction (Z-score of average ΔGE values) for 47-gene viral mimicry signature, and (B) Heatmap depicts the relative differential induction of genes used for the viral mimicry signature in Fun-CIM and Dys-CIM tumors in the NKI/SMC cohorts using Z-score of average ΔGE values. (C–D) Corresponding analysis of the viral mimicry signature in the NEO cohort, showing the overall program induction (C) and individual signature genes (D). (E-F) Viral mimicry induction in TNBC cell lines following epirubicin treatment, showing the overall program induction (E) and individual signature genes (F). (G–L) Boxplots show differential gene induction (Δlog2[TPM+1]) in Dys-CIM and Fun-CIM cell lines for key cytosolic nucleic acid sensing genes, including *IFIH1* (G), *CGAS* (H), *STING1* (I), *AIM2* (J), *DHX58* (K), and *DDX58* (RIG-I) (L). Adjacent stacked bar plots show the proportion of samples exhibiting positive induction (ΔGE > 0; green) versus no or negative induction (ΔGE ≤ 0; red) within each CIM trajectory. Fun-CIM cell lines exhibited greater induction of multiple cytosolic nucleic acid sensors, including *IFIH1, STING1, AIM2,* and *DHX58*. p values above boxplots in (G–L) indicate comparisons of ΔGE between CIM trajectories using the Wilcoxon rank-sum test, whereas p values above stacked bar plots indicate comparisons of the proportions of samples exhibiting positive versus no/negative induction using Fisher’s exact or chi-squared test, as indicated. FDR-adjusted p values for viral mimicry program and gene-level comparisons in (A–F) are indicated as follows: * ≤ 0.05; ** ≤ 0.01; *** ≤ 0.005; **** ≤ 0.001.

**Supplemental Figure S16.**
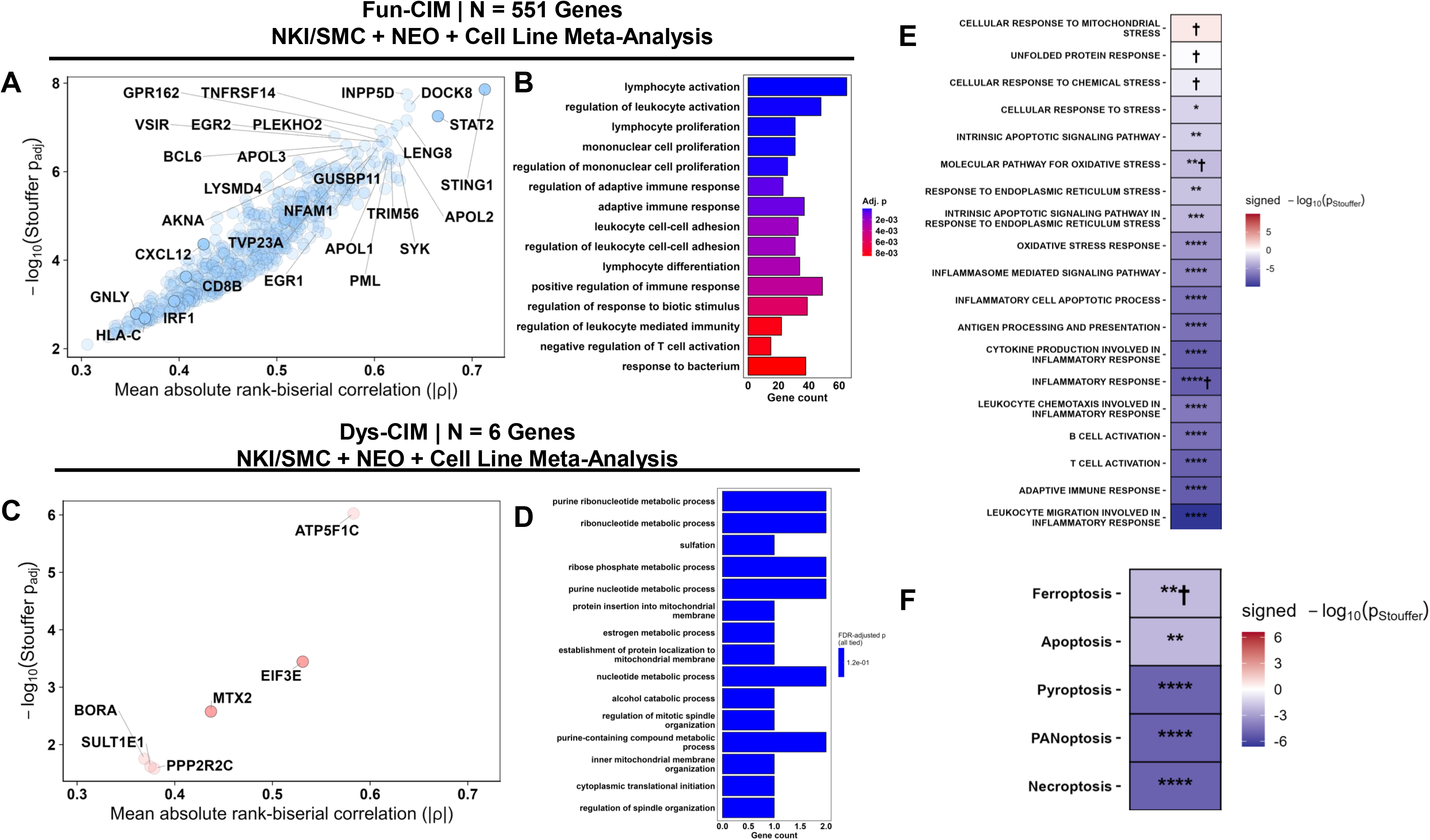
Conserved Fun-CIM and Dys-CIM transcriptional programs across patient cohorts and TNBC cell lines. (A, C) Stouffer conservation analysis of directionally concordant genes across NKI/SMC, NEO, and TNBC cell lines identified 551 conserved Fun-CIM (A) and 6 Dys-CIM (C) genes. (B, D) Gene Ontology overrepresentation analysis showed enrichment of immune and inflammatory programs in Fun-CIM (B) and protein maintenance and metabolic programs in Dys-CIM (D). (E–F) Conservation analysis of CIM-related (E) and regulated cell-death (F) pathways demonstrated conserved induction of immunostimulatory and inflammatory cell-death programs in Fun-CIM across all three datasets.

**Supplemental Figure S17.**
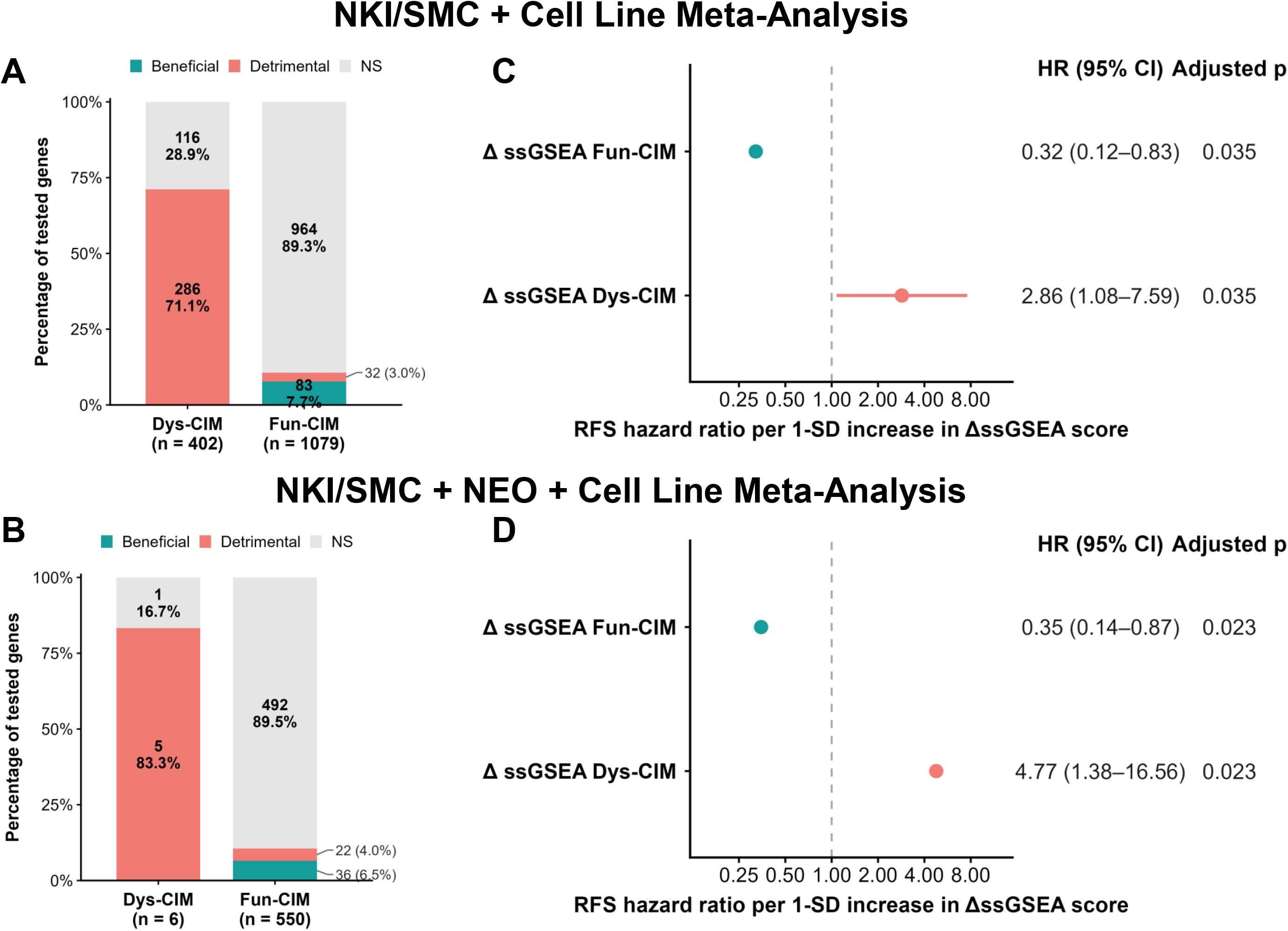
Association of conserved Fun-CIM and Dys-CIM transcriptional remodeling with recurrence-free survival in the NKI cohort. (A–B) Directional distribution of gene-level associations with recurrence-free survival (RFS) for conserved Dys-CIM and Fun-CIM gene sets identified by the NKI/SMC + TNBC cell line (A) and NKI/SMC + NEO + TNBC cell line (B) meta-analyses. For each gene, change was calculated as surgery minus biopsy Δlog₂(TPM+1) expression and standardized within gene. Genes were evaluated individually using univariable Cox proportional-hazards models and classified as beneficial (*P <* 0.05, HR < 1), detrimental (*P <* 0.05, HR > 1), or not significant (NS). (C–D) Associations between longitudinal changes in ssGSEA scores for the corresponding conserved Fun-CIM and Dys-CIM signatures and RFS. Hazard ratios represent the association per 1-SD increase in ΔssGSEA score. Increased Fun-CIM scores were associated with improved RFS, whereas increased Dys-CIM scores were associated with poorer RFS. HR, hazard ratio; CI, confidence interval; ssGSEA, single-sample gene set enrichment analysis.

**Supplemental Figure S18.**
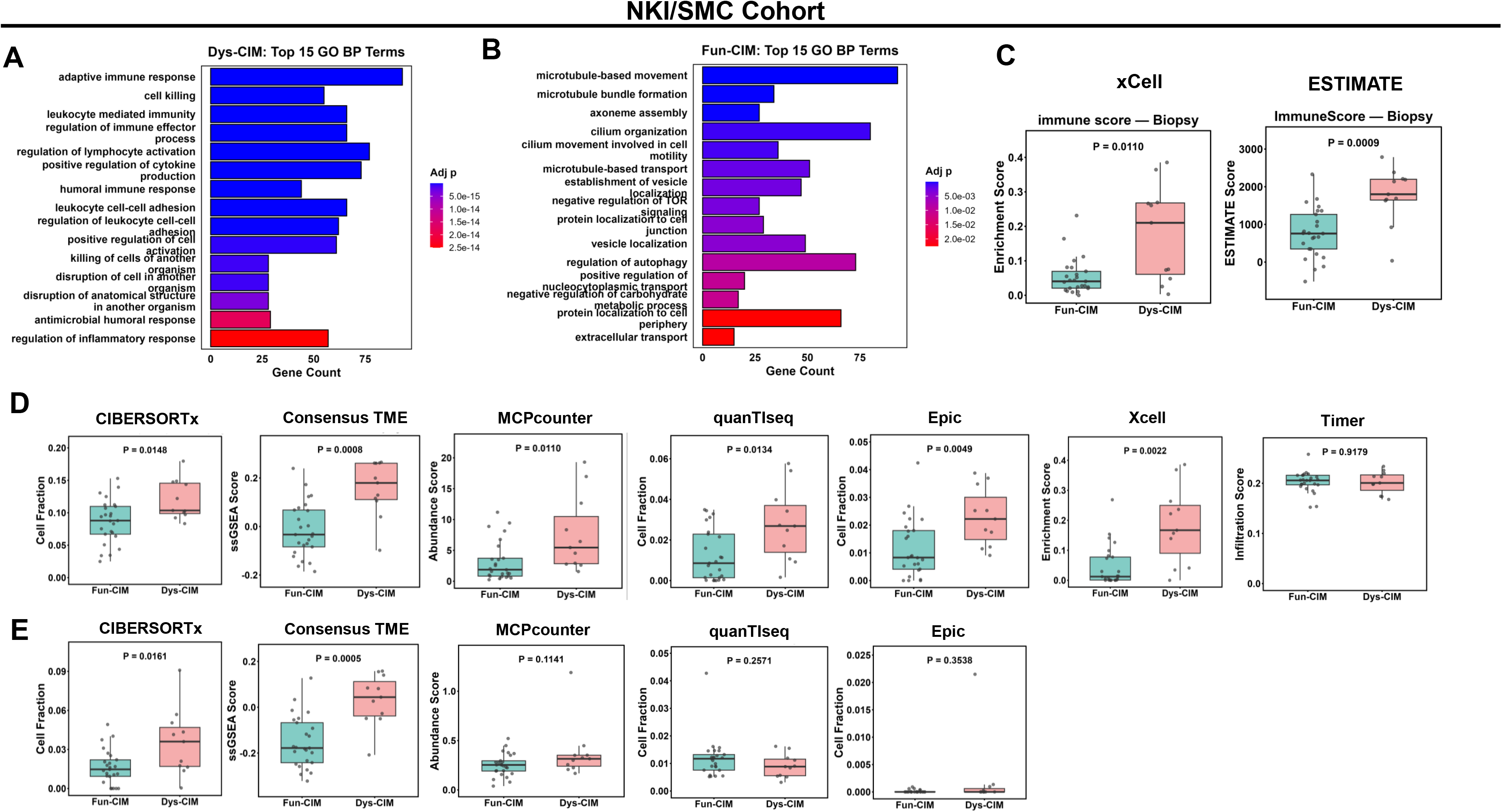
Dys-CIM tumors in the NKI/SMC cohorts exhibit baseline overrepresentation of immune-related transcriptional programs and greater immune enrichment than Fun-CIM tumors. (A–B) Top 15 enriched Gene Ontology Biological Process (GO BP) terms among genes with higher baseline expression in Dys-CIM (A) or Fun-CIM (B) tumors from the combined NKI/SMC cohort. Baseline differentially expressed genes were identified using limma-trend with Benjamini–Hochberg-adjusted p ≤ 0.05 and |log₂ fold change| ≥ log₂(1.25). (C) Baseline overall immune enrichment in Fun-CIM (*n* = 25) and Dys-CIM (*n* = 11) tumors, estimated using the xCell and ESTIMATE immune scores. (D–E) Baseline abundance of CD8⁺ T cells (D) and NK cells (E) estimated using the indicated immune-deconvolution methods. Dys-CIM tumors exhibited higher baseline CD8⁺ T-cell abundance across multiple methods, whereas differences in NK-cell abundance were less consistent. Displayed p *value*s were calculated using two-sided Wilcoxon rank-sum test.

**Supplemental Figure S19.**
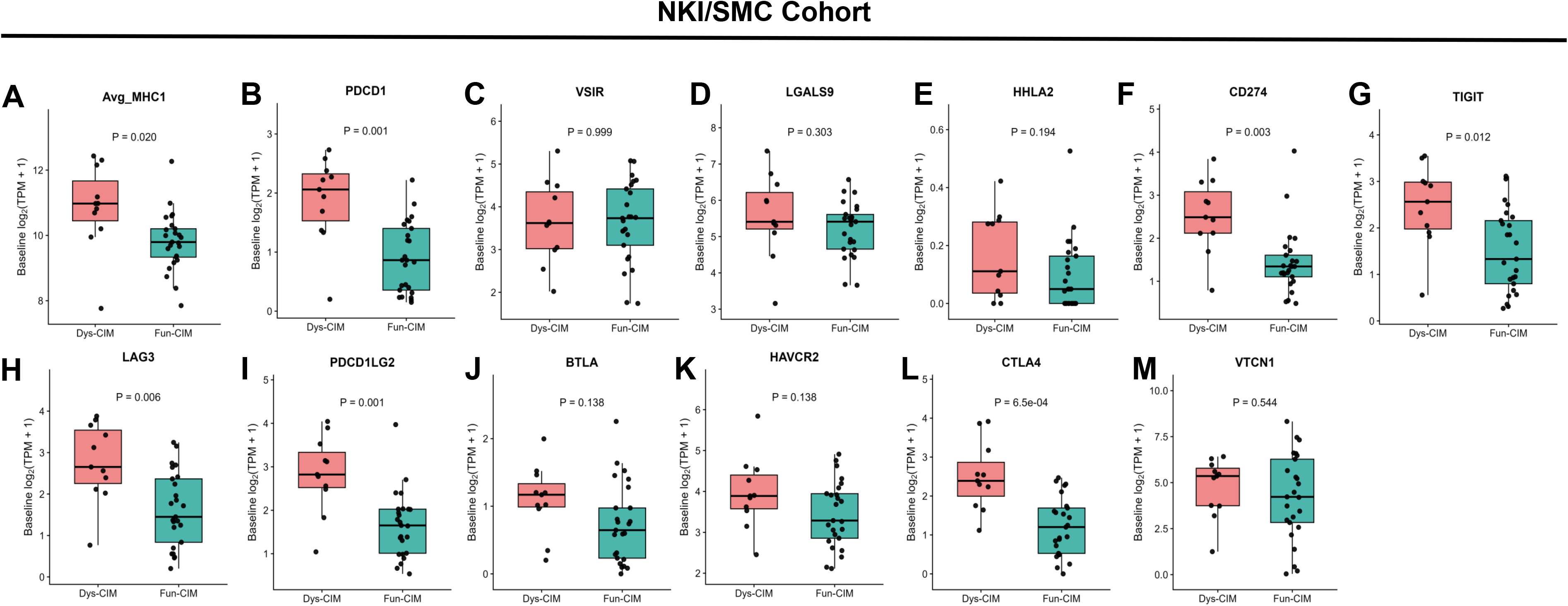
Dys-CIM tumors exhibit higher baseline MHC-I and immune checkpoint-related gene expression than Fun-CIM tumors. (A–M) Boxplots show baseline log₂(TPM + 1) expression of Avg_MHC1 (mean expression of *HLA-A, HLA-B,* and *HLA-C*) (A) and immune checkpoint-related genes, including *PDCD1* (B), *VSIR* (C), *LGALS9* (D), *HHLA2* (E), *CD274* (PD-L1) (F), *TIGIT* (G), *LAG3* (H), *PDCD1LG2* (PD-L2) (I), *BTLA* (J), *HAVCR2* (K), *CTLA4* (L), and *VTCN1* (M) in Dys-CIM (n = 11) and Fun-CIM (n = 25) tumors from the NKI/SMC cohorts. Dys-CIM tumors exhibited significantly higher baseline Avg_MHC1 and expression of multiple immune checkpoint-related genes, including *PDCD1, LGALS9, CD274, TIGIT, LAG3, PDCD1LG2,* and *CTLA4*. Displayed p values are Benjamini–Hochberg FDR-adjusted and were estimated using limma with robust, trend-aware empirical Bayes moderation.

## Notes

https://github.com/Gbadamosi-Lab/CIMIC

### Competing Interest Statement

The authors have declared no competing interest.

