## Supplemental Methods for "Unsupervised transcriptomic analysis of paired pre- and post-treatment specimens reveals divergent chemoimmunomodulatory induction trajectories in breast cancer"

Department of Pharmacotherapy and Translational Research, College of Pharmacy, University of Florida, 1345 Center Drive, Room PG-05B Gainesville, FL 32610

Running Title: Chemoimmunomodulation Induction Trajectories Pipeline

### Supplemental Methods

#### SM1. RNA-Sequencing Datasets Harmonization

RNA-seq expression data from the NKI, SMC, and NEO cohorts were harmonized prior to combining the datasets. Technical replicates within NKI were averaged at the raw TPM level prior to log transformation (log_2_(TPM+1)), whereas the SMC and NEO expression matrices, which contained single measurements per sample were directly log transformed. Gene identifiers were updated to reflect the most current HGNC-approved gene symbols using the HUGO Gene Nomenclature Committee (HGNC) complete gene set with predefined rules to resolve multiple identifiers mapping to the same HGNC symbol (1). Specifically, when multiple source identifiers mapped to the same approved symbol and the approved symbol was present among the original identifiers, the original approved-symbol entry was retained and any aliases were removed. In addition, when multiple distinct source identifiers mapped to the same approved symbol without an original identifier corresponding to that symbol, all entries were excluded to avoid ambiguous gene assignment. The resulting gene mappings were validated to ensure one-to-one correspondence between source genes and standardized symbols. Post-harmonization, data from the NKI and SMC cohorts were combined to form the first dataset. Data from the NEO cohort was analyzed independently.

#### SM2. CIMIC Chemoimmunomodulation Pathway Selection Criteria and Filtering

To comprehensively map the multidimensional landscape of chemoimmunomodulation (CIM), we performed a systematic curation of gene sets from the Molecular Signatures Database using the msigdbr R package (v 25.1.1) (2). Candidate pathways were identified through targeted keyword searches designed to capture the intersection of tumor-intrinsic stress adaptation and extrinsic immunomodulatory signaling, specifically utilizing the terms: "Inflammatory," "Inflammation," "Immune Response," "Stress," "Antigen Presentation," and "Pattern AND Recognition." To focus on well-established pathways supported by strong evidence, the search space was restricted to three primary MSigDB collections: (i) Hallmark (H) gene sets, prioritized for their refined and non-redundant representation of coherent biological states; (ii) Curated (C2) gene sets, restricted specifically to Canonical Pathways (CP)**,** including KEGG, Reactome, and WikiPathways; and (iii) Ontology (C5) gene sets. The search was further limited to the Biological Process (BP) sub-collection to emphasize functional induction over cellular localization. Following initial retrieval, gene sets were cross-referenced against established literature to ensure their documented involvement in chemotherapy-induced remodeling of the tumor microenvironment. This rigorous selection process resulted in a finalized compendium of 19 CIM-related pathways, encompassing 3,189 unique genes, served as the feature input for the CIMIC pipeline. Minimal biological redundancy was confirmed via Jaccard Index analysis (max value: 0.381, median value: 0.009, mean value: 0.026).

To focus on tumor-intrinsic and broad immune signaling rather than clonal lymphocyte expansion, we implemented a rigorous filtering step to remove highly variable immune receptor components. We excluded genes with prefixes associated with B-cell immunoglobulin heavy and light chains (e.g., *IGHV, IGHD, IGLV, IGKV*) and T-cell receptor (TCR) alpha, beta, gamma, and delta chains (e.g., *TRAV, TRBV, TRAC*). Thus, ensuring that the CIMIC classifications reflect generalized immunomodulatory states rather than dominant lymphocyte clones.

#### SM3. CIMIC Algorithmic and Pipeline Implementation Details

For all PaCMAP (v0.8.0) dimensionality reduction analyses, the package default parameters were used including: n_components = 2, n_neighbors = 10, MN_ratio = 0.5, FP_ratio = 2.0, metric = “euclidean”, and init = “pca”. For ConsensusClusterPlus (v1.73.0) the parameters were as follows: hierarchical clustering (hc) was used as the base clustering algorithm, Ward's linkage (D2) for inner and final linkage, Euclidean distance as the distance metric, 5,000 as the default number of resampling iteration and pFeature was set to 1.0. The item-resampling proportion (pItem) was selected adaptively as described below. To avoid undefined item-consensus values that can arise at low item-sampling fractions, clustering began at pItem = 0.80 and was incremented in steps of 0.05 (to a maximum of 1.0) until the resulting item-consensus matrix, computed via calcICL, contained no missing values. If no candidate pItem value in this range eliminated all missing values, the maximum candidate number of clusters (*k*) was reduced to the largest *k* for which item-consensus was fully defined across the pItem sweep, and the adaptive procedure was repeated at that reduced *k*. When the maximum stable *k* resolved to 2, ConsensusClusterPlus was run with maximum *k* = 3 and only the *k* = 2 partition was retained, as ConsensusClusterPlus requires maximum *k* ≥ 3 to compute a well-defined item-consensus table at *k* = 2.

The optimal number of clusters (*k)*, used for consensus clustering procedures in the CIMIC pipeline, is determined using a composite ranking framework that integrates measures of cluster separation and stability metrics across the tested range of *k* values. Cluster separation is evaluated using the average of the silhouette coefficients (*s_avg_*) defined as the mean of the mean silhouette widths calculated in the PaCMAP embedding space and the original ΔGE space (3), while cluster stability is assessed using the proportion of ambiguous clustering (PAC) (4), which is defined as the empirical CDF difference F(0.9) − F(0.1) over the off-diagonal consensus matrix entries and quantifies instability in pairwise sample co-clustering across resampling iterations, and the cluster consensus score (CCS) (5), which quantifies the overall reproducibility of cluster structure. Evaluation metric values for each *k* are independently ranked across the tested range of *k* values, such that higher separation and stability correspond to lower rank values (higher s_avg_ and CCS and lower PAC). The overall rank for each *k* is calculated as the unweighted sum of ranks across all evaluation metrics, and the optimal *k* is defined as the *k* with the lowest sum. In the event of a tie, CCS was used as the default tie-breaking metric; the tie-breaking metric can be specified by the user. For all consensus clustering procedures, values of *k* = 2 to *k* = 5 were evaluated to capture parsimonious population-level structure while avoiding overpartitioning of the relatively small patient cohorts.

For limma-based linear models with empirical-Bayes variance moderation, models were fit using limma::lmFit followed by empirical-Bayes moderation with eBayes (robust = TRUE, trend = TRUE) using a design matrix of ~ cluster. Two-cluster solutions are evaluated using moderated *t*-statistics to identify genes with differential mean ΔGE between clusters, whereas multi-cluster solutions are evaluated using moderated *F*-statistics to identify genes with differential mean ΔGE across clusters. For iterative-stability guided refinements, a maximum of 50 iterations is permitted per procedure. Convergence is typically achieved within 3-5 iterations.

Because the clustering step operates on an iteratively refined gene subset, two instability conditions were handled explicitly: (i) if PaCMAP embedding failed for a given feature subset, the most recently converged (prior-iteration) feature set was retained and refinement halted; (ii) if iterative refinement immediately converged to zero genes surviving limma-based gene filtering the top 10% of tested genes ranked by adjusted p-value then effect size were retained as a fallback set to permit continued clustering.

#### SM4. Stability and Reproducibility of CIMIC

To confirm the stability and reproducibility of CIMIC, and its robustness to PaCMAP parameterization and stochastic variation, we analyzed CIMIC cluster assignments across multiple PaCMAP dimensional representations (n_components = 2, 5, 10, and 15) (6) using 15 random seeds per n_component. Concordance of cluster assignments across analyses was quantified using the adjusted Rand index (ARI) in NKI/SMC and NEO (7). In NKI/SMC, the two-cluster solution reported and described in the main text demonstrated complete concordance across all evaluated PaCMAP parameterizations and random seeds (mean ARI = 1.00; SD = 0) and were highly selected across runs (**SM Table 1**), whereas the three-cluster solution (described below) were less selected across runs but still demonstrated high concordance (mean ARI = 0.914; SD = 0.040; range= 0.886-0.942). Similarly, the two-cluster solution in NEO demonstrated high concordance across PaCMAP parameterizations and random seeds (mean ARI = 0.972; SD = 0.072; range = 0.789–1.00). The two-cluster solution was highly selected across all seeds.

| **SM Table 1. Adjusted Rand Index Analysis of CIMIC Solutions** | | | | | | | |
| --- | --- | --- | --- | --- | --- | --- | --- |
| **NKI/SMC** | **n_components** | **k** | **ari_mean** | **ari_sd** | **ari_min** | **ari_max** | **ari_n** |
|  | **2** | 2 | 1 | 0 | 1 | 1 | 12 |
|  |  | 3 | 0.886 | 0.046 | 0.832 | 0.913 | 3 |
|  | **5** | 2 | 1 | 0 | 1 | 1 | 12 |
|  |  | 3 | 0.942 | 0.05 | 0.913 | 1 | 3 |
|  | **10** | 2 | 1 | 0 | 1 | 1 | 14 |
|  | **15** | 2 | 1 | 0 | 1 | 1 | 15 |
| **NEO** | **2** | 2 | 0.970 | 0.074 | 0.789 | 1 | 14 |
|  |  | 3 | NA | NA | NA | NA | 1 |
|  | **5** | 2 | 0.972 | 0.072 | 0.789 | 1 | 15 |
|  | **10** | 2 | 0.972 | 0.072 | 0.789 | 1 | 15 |
|  | **15** | 2 | 0.972 | 0.072 | 0.789 | 1 | 15 |

#### SM5. Correlation Between ∆GE and Pre-treatment GE

To answer the question of whether CIM gene induction values [∆(log_2_(TPM+1))] values were merely a function of pre-treatment CIM gene expression, we performed Spearman's correlation analysis between pre-treatment gene expression and gene induction, restricting the profiles to CIM genes (**Figure SM1**). We similarly performed correlation analysis between pre-treatment CIM gene profile and post-treatment CIM profile as a control, given that the gene-expression profile from the same patient pre-treatment and post-treatment should remain similar because they share stable, patient-specific features, partially conserved tissue composition, and technical factors. Our results demonstrate that CIM profiles had weak correlations comparing pre-treatment to gene induction values. In contrast, the internal positive control correlations, comparing pre-treatment CIM gene expression and post-treatment values, tended to have strong correlations.

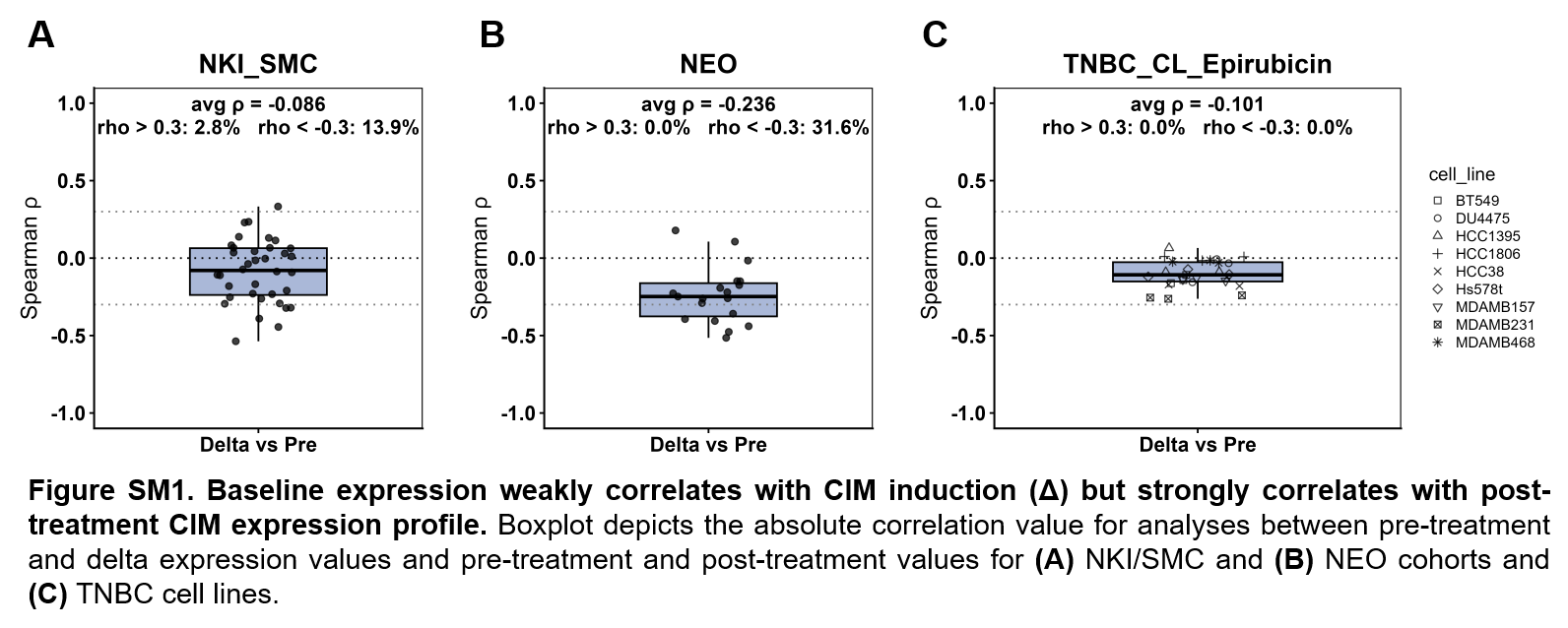

#### SM6. Batch Effect Analyses

To assess whether cohort identity contributed substantially to variation in the delta-expression space used by CIMIC, we computed Δlog2(TPM+1) per patient, centered and scaled the matrix, and performed PCA analysis on the induction values (**Figure SM2**). In the PC1–PC2 projection (17.2% and 11.1% variance), samples from the Netherlands Cancer Institute and Samsung Medical Center cohorts were well intermixed, with no visual separation by cohort, thus indicating minimal batch effect between both cohorts.

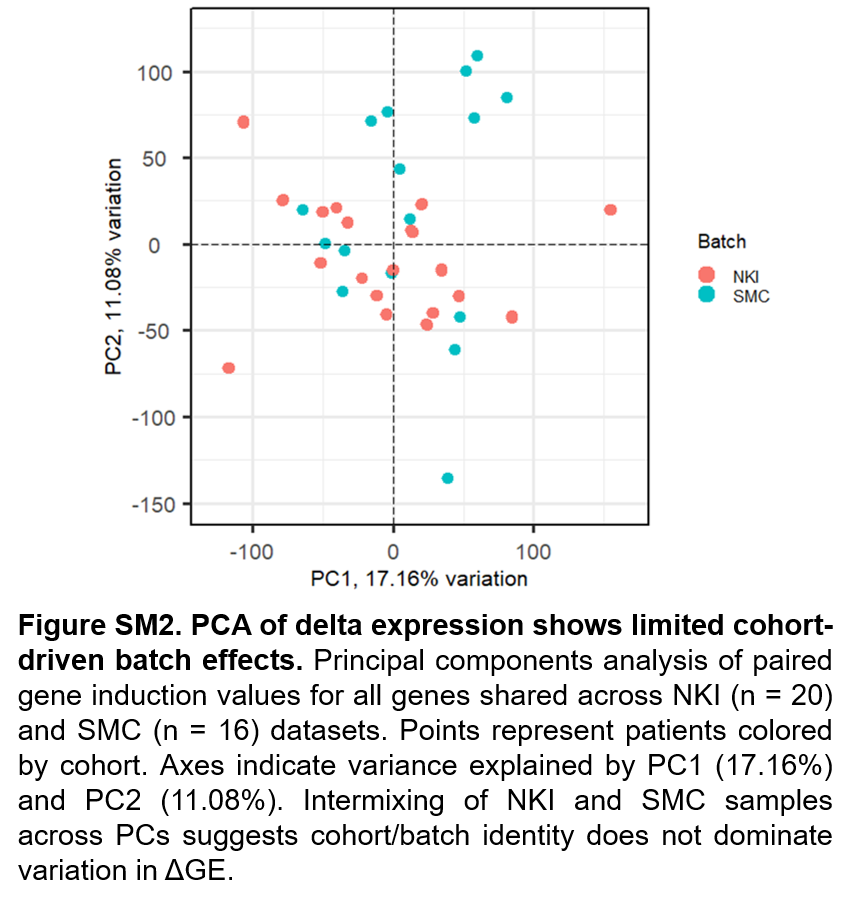

#### SM7. A note on model selection and alternative clustering solutions

In addition to the selected optimal solution at k = 2, k = 3 also yielded a strong biologically plausible clustering solution, highlighting the flexibility of the CIMIC pipeline resolve CIM trajectories at different levels of granularity (**Figure SM4A-SMA4B**). In this solution, the Fun-CIM cluster subdivided into two related subclusters, one characterized by broad immune-activation with increased cytotoxic immune cell immune abundance, and one
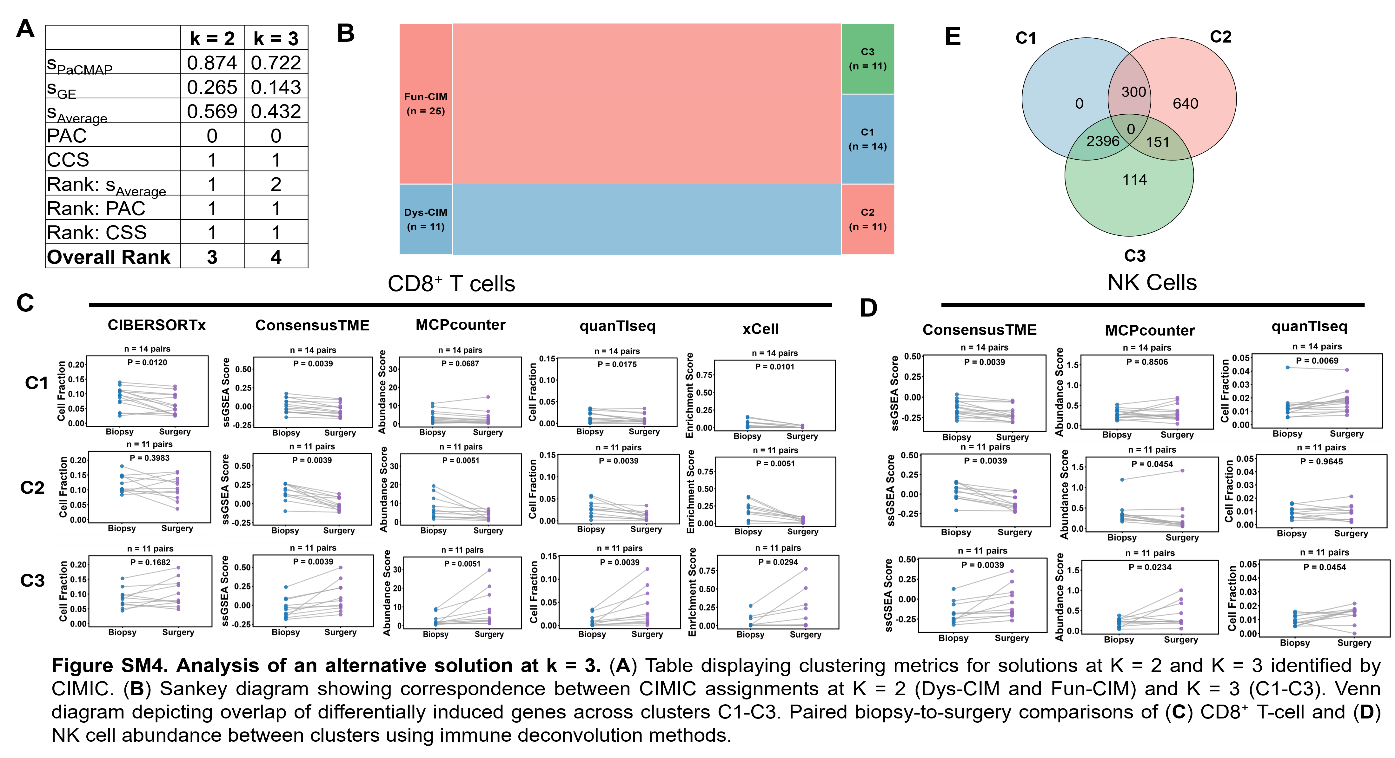
 characterized by immune-activation without limited increases in cytotoxic immune cell abundance (**Figure SM4C-SM4D**). Although this solution provided additional resolution, the K = 2 solution was maintained for primary analyses, given the large overlap in the CIM genes induced between subdivisions of the Fun-CIM trajectory (**Figure SM4E**). Future work will aim to analyze the K = 3 solution to characterize features along the continuity of CIM.
